# The HAPPE plus Event-Related (HAPPE+ER) Software: A Standardized Processing Pipeline for Event-Related Potential Analyses

**DOI:** 10.1101/2021.07.02.450946

**Authors:** A.D. Monachino, K.L. Lopez, L.J. Pierce, L.J. Gabard-Durnam

## Abstract

Event-Related Potential (ERP) designs are a common method for interrogating neurocognitive function with electroencephalography (EEG). However, the gold standard of preprocessing ERP data is manual-editing – a subjective, time-consuming processes. A number of automated pipelines have recently been created to address the need for standardization, automation, and quantification of EEG data processing; however, few are optimized for ERP analyses (especially in developmental or clinical populations). To fill this need, we propose and validate the HAPPE plus Event-Related (HAPPE+ER) software, a standardized and automated processing pipeline optimized for ERP analyses in EEG data. HAPPE+ER processes event-related potential data from raw files through a series of filtering, line noise reduction, bad channel detection, artifact rejection from continuous data, segmentation, and bad segment rejection methods. HAPPE+ER also includes post-processing reports of both data quality and pipeline quality metrics to facilitate the evaluation and reporting of data processing in a standardized manner. Finally, HAPPE+ER includes a post-processing script to facilitate generating ERP figures and measures for statistical analysis. We describe multiple approaches with both adult and developmental data to optimize and validate pipeline performance. HAPPE+ER software is freely available under the terms of GNU General Public License at https://github.com/PINE-Lab/HAPPE.

## Introduction

There is growing momentum to standardize and automate electroencephalography (EEG) and event-related potential (ERP) processing to meet the needs of contemporary electrophysiological studies. Until recently, the gold standard of preparing EEG/ERP data for analysis involved removing artifact-laden segments through subjective manual editing. However, this process can result in significant data loss, especially in data from developmental and clinical populations characterized by high levels of artifact. This process has also become difficult to scale as sample sizes and electrode densities for recording have both increased substantially over the last decade. Furthermore, the subjective nature of manual editing impedes comparisons across EEG acquisition systems, datasets, and laboratories. The solution appears in automated, standardized processing. However, early processing software was often limited to single stages of EEG processing (Haresign et al., 2021; Lawhern et al., 2013; Leach et al., 2020; Mognon et al., 2011; Nolan et al., 2010; Winkler et al., 2011), developed only on data with low levels of artifact, and lacked imbedded metrics to quantitatively assess their performance. HAPPE software (Gabard-Durnam et al., 2018) proposed a solution to these limitations by providing an automated, quantifiable, and standardized method of processing EEG data that is effective with high levels of artifact as seen in developmental and clinical populations.

HAPPE is not alone in its endeavors; a growing number of pipelines, scripts, and software now also address the need for standardized pre-processing methods for EEG data. With the breadth of available tools comes a diversity of approaches to EEG data processing (Bigdely-Shamlo et al., 2015; Cassani et al., 2017; da Cruz et al., 2018; Debnath et al., 2020; Desjardins et al., 2021; Gabard-Durnam et al., 2018; Gramfort et al., 2014; Hatz et al., 2015; Leach et al., 2020; Oostenveld et al., 2011; Pedroni et al., 2019; Tadel et al., 2011). Few empirical comparisons have been made between pipelines, and it may be difficult for researchers to assess which pipeline works best on their EEG data. Moreover, pipelines differ in their limitations on the kinds of analyses that can be performed post-processing. Some pipelines are restricted to preparing data for time-frequency analyses or resting-state EEG data (Cassani et al., 2017; Gabard-Durnam et al., 2018; Hatz et al., 2015). Pipelines also differ in the populations for which they have been validated. Namely, the majority of pipelines are tested and validated using data from healthy adults (Bigdely-Shamlo et al., 2015; Cassani et al., 2017; da Cruz et al., 2018; Mognon et al., 2011; Nolan et al., 2010; Pedroni et al., 2019). Far fewer specify compatibility and validation with developmental or clinical populations, whose data often have physiological and acquisition differences from healthy adult data. Exceptions include MADE (Debnath et al., 2020)(validated in developmental data); EEG-IP-L (Desjardins et al., 2021), and HAPPE (Gabard-Durnam et al., 2018)(validated in clinical and developmental data). The optimal pipeline would offer validated solutions suitable to developmental, clinical, and adult EEG/ERP processing needs to facilitate comparisons across studies and ages.

Here we propose and validate the HAPPE plus ER (HAPPE+ER) software to address these limitations for ERP analyses, improve on the original HAPPE pre-processing strategies, and increase accessibility across acquisition setups and user coding fluencies. This software includes both code to process ERP data and code that enables the efficient, automated creation of post-processing ERP figures and measures, including (but not limited to) peak amplitudes, latencies, and area under the curve. HAPPE+ER (pronounced “happier”) incorporates pre- processing steps for ERP designs that we validate in adult, developmental, and clinical data.

HAPPE+ER also introduces a novel pipeline quality report of metrics reflecting each pre- processing step’s performance on ERP data to aid researchers in assessing whether the pipeline is effectively pre-processing their data. That is, HAPPE+ER now includes complementary, quantifiable measures of quality for both data inputs (data quality report) and processing methods (pipeline quality report). In conjunction with these changes, data can be input from an increased array of file formats and acquisition layouts, including nets from EGI, BioSemi, and Brain Products. These changes greatly expand the breadth of data for which HAPPE+ER is suitable while maintaining standardization across a variety of input parameters and output analyses, rendering HAPPE+ER a flexible software suited to electrophysiological processing across the lifespan and populations. The following sections detail and justify the HAPPE+ER pipeline’s processing steps for ERP analyses, compare multiple approaches to artifact rejection in ERP data, outline post-processing report metrics, and demonstrate HAPPE+ER’s effectiveness with adult and developmental ERP datasets.

### HAPPE+ER Pipeline Steps

### HAPPE+ER Data Inputs

HAPPE+ER accommodates multiple types of EEG files with different acquisition layouts as inputs, with additional options on top of those previously supported in HAPPE 1.0 software.

See Table 1 for complete layout and formatting options accepted by the HAPPE+ER pipeline. For .set formatted files, the correct channel locations should be pre-set and embedded in the file (e.g., by loading it into EEGLAB (Delorme and Makeig, 2004) and confirming the correct locations) prior to running through HAPPE+ER. Each batch run of HAPPE+ER must include files collected with the same channel layout (company, net type, and electrode number) and paradigm (resting-state or event-related), each of which users must specify for a given run. The same is true of file formats, in that a single run will support only a single file type across files, specified by the user. HAPPE+ER processes data collected with any sampling rate, and files within a single run may differ in their individual sampling rates.

**Table 1.** A list of acquisition layouts and associated file types supported by HAPPE+ER. Includes company name and number of channels/leads.

| Company | Net Name | Number of Channels | File Types Supported |
| --- | --- | --- | --- |
| Electrical Geodesics, Inc. | Geodesic Sensor Net | 64 | .mat, .raw, .set, .mff |
| Electrical Geodesics, Inc. | Hydrocel Geodesic Sensor Net | 32, 64, 128, 256 | .mat, .raw, .set, .mff |
| BioSemi | BioSemi Headcap | 16, 64 | .set |
| BioSemi | BioSemi Headcap | 32, 128 | .mat, .set |
| BrainProducts | Standard BrainCap | 22, 32, 64, 96, 128 | .set |
| BrainProducts | Wet-Sponge R-Net for actiCHamp Plus | 32, 64, 96, 128 | .set |
| Neuroscan | Quik-Cap | 32, 64, 128, 256 | .cdt, .set |
| Low-Density Layouts |  | 30 channels or less | .mat, .set |

### Specifying EEG Channels for Inclusion/Exclusion

HAPPE+ER offers a variety of options for channel selection such that the user can choose the channels that best fit the needs of their dataset. HAPPE+ER does not restrict the user to a specific number of channels, as no later processing steps rely on channel number for robust processing. Specifically, HAPPE+ER supports the following options: 1) Selecting all channels, which results in each channel included in the dataset being included in the following processing steps. 2) Alternatively, the user can select a subset of channels of interest, which can be selected via an inclusion or exclusion method. Either method of selecting a channel subset will automatically include the 10-20 system electrodes appropriate for the channel layout.

Selecting the option to include user-specified channels will remove every channel not included in the user-specified list (except for the 10-20 channels that are automatically included) from subsequent processing with the inability to recover them later. For example, for data from a 128-channel net where the user selects 20 channels, the post-HAPPE+ER processed data will contain only data for those 20 selected channels plus the 19 10-20 channels for a total of 39 channels.

Selecting the option to exclude channels does the opposite, excluding all user-specified channels and keeping those not included in the user-specified list. For example, for data from a 128-channel net where the user specifies 8 channels, the post HAPPE+ER processed data will contain the remaining 120 channels. These options increase the ease of selecting channels based on the number of channels a user is interested in examining.

### Electrical (Line) Noise Removal

HAPPE+ER removes electrical noise (e.g., 60 or 50 Hz artifact signal) from EEG through the multi-taper regression approach implemented by the CleanLine program (Mullen, 2012).

Multi-taper regression can remove electrical noise without sacrificing or distorting the underlying EEG signal in the nearby frequencies, drawbacks of the notch-filtering approach to line-noise removal (Mitra and Pesaran, 1999). Specifically, HAPPE+ER applies the updated version of CleanLine’s multi-taper regression which is more effective at removing line noise than the original version present in HAPPE 1.0. The legacy CleanLine version from HAPPE 1.0 is available as an option to the user, however the updated version is registered as the default.

CleanLine’s multi-taper regression scans for line-noise signal near the user-specified frequency ± 2 Hz, a 4-s window with a 1-s step size and a smoothing tau of 100 during the fast Fourier transform, and a significance threshold of p = 0.01 for sinusoid regression coefficients during electrical noise removal. This process is highly specific to the frequency of electrical noise, which the user can specify to be 60 Hz or 50 Hz. Pipeline quality control metrics for the effectiveness of line noise removal are automatically generated in HAPPE+ER and discussed in detail as part of the subsequent “Quality Control Metrics” section of this manuscript.

### Filtering

HAPPE+ER applies an automatic low-pass filter at 100 Hz prior to artifact rejection. Additional filtering for ERP analyses as determined by the user occurs after artifact rejection. Note this filtering strategy differs from that in HAPPE 1.0, where a 1Hz high-pass filter was applied to all files to facilitate optimal ICA decomposition (Winkler et al., 2015). HAPPE 1.0 precluded ERP analyses since filtering at 1Hz excludes frequencies of interest in ERP analyses.

### Bad Channel Rejection (Optional)

HAPPE+ER can detect and remove channels that do not contribute to usable brain data due to high impedances, damage to the electrodes, insufficient scalp contact, and excessive movement or electromyographic (EMG) artifact throughout the recording.

HAPPE+ER performs bad channel detection that is suitable for ERP data and expands the classes of bad channels that can be detected relative to prior automated pipeline options by combining EEGLAB’s Clean Rawdata functions with power spectral evaluation steps as follows. First, HAPPE+ER runs the Clean Rawdata ‘Flatline Criterion’ to detect channels with flat recording lengths longer than 5 seconds, indicating no data collected at that location. If the channel contains a flatline that lasts longer than 5 seconds, the channel is marked as bad. After flat channels have been removed, HAPPE+ER includes a spectrum-based bad channel detection step similar to that used in HAPPE 1.0 where bad channel detection is achieved by twice evaluating the normed joint probability of the average log power from 1 to 125 Hz across the user-specified subset of included channels. Channels whose probability falls more than 3 standard deviations from the mean are removed as bad channels. While the HAPPE 1.0 method of legacy detection proved to be suboptimal for our test dataset (see Table 2), evaluating the joint probability of average log power from 1 to 100Hz was useful for optimizing bad channel detection alongside Clean Rawdata. HAPPE+ER thus includes a spectrum evaluation step with thresholds of -5 and 3.5 standard deviations. Here, the standard deviations are not symmetric as artifact in the EEG signal mostly produces power spectrums with positive standard deviations from the mean. Channels near the reference electrode will have reduced amplitudes by virtue of sharing variance with the reference, rather than due to artifact, but will score below the mean in their average log power accordingly. To avoid falsely rejecting (good) channels near the reference electrode(s), the negative standard deviation threshold is set liberally at –5 standard deviations. Finally, HAPPE+ER uses Clean Rawdata’s ‘Line Noise Criterion’ and ‘Channel Criterion’ to detect additional bad channel cases. The line noise criterion identifies whether a channel has more line noise relative to neural signal than a predetermined value in standard deviations based on the total channel population, set in HAPPE+ER to 6 standard deviations. Channel criterion sets the minimally acceptable correlation value between the channel in question and all other channels. If a channel’s average correlation is less than the preset value, it is considered abnormal and marked as bad. HAPPE+ER uses a correlation threshold of .8 to identify bad channels through this approach.

**Table 2.** Performance of bad channel parameters tested on twenty files from an EGI dataset. The settings for each individual step are separated by a newline within the cell.

| Bad Channel Parameters | Accuracy<br>(780 Total Channels) | False Positive<br>(741 Good Channels) | False Negatives<br>(39 Bad Channels) |
| --- | --- | --- | --- |
| HAPPE 1.0 Legacy Detection | 92.4% | 41 | 18 |
| Flatline = 5; Channel Corr = .1; Line Noise Ratio = 20<br>Spectrum = -5, 3.5<br>Channel Corr = .8; Line Noise Ratio = 6 | 97.6% | 9 | 10 |
| Flatline = 5; Channel Corr = .1; Line Noise Ratio = 20<br>Spectrum = -5, 3.5<br>Channel Corr = .75; Line Noise Ratio = 6 | 97.2% | 7 | 15 |
| Flatline = 5; Channel Corr = .1; Line Noise Ratio = 20<br>Spectrum = -5, 3.5<br>Channel Corr = .8; Line Noise Ratio = 5 | 97.3% | 11 | 10 |
| Flatline = 5; Channel Corr = .1; Line Noise Ratio = 20<br>Spectrum = -4, 3.5<br>Channel Corr = .8; Line Noise Ratio = 6 | 97.2% | 12 | 10 |
| Flatline = 5; Channel Corr = .1; Line Noise Ratio = 20<br>Spectrum = -6, 3.5<br>Channel Corr = .8; Line Noise Ratio = 6 | 97.6% | 9 | 10 |

**Table 3.** Average percent segments retained and correlation values pre- and post-waveleting in adult, 4-month, and 10-month data for wavelet level optimization.

| Processing Strategy | Average Percent Segments Retained | r pre/post wavelet |  |  |  |  |  |  |  |  |  |  |
| --- | --- | --- | --- | --- | --- | --- | --- | --- | --- | --- | --- | --- |
|  |  | all data | 0.5 Hz | 1 Hz | 2 Hz | 5 Hz | 8 Hz | 12 Hz | 20 Hz | 30 Hz | 45 Hz | 70 Hz |
| Adult data |  |  |  |  |  |  |  |  |  |  |  |  |
| Hand-Edit | 86.08 | NA | NA | NA | NA | NA | NA | NA | NA | NA | NA | NA |
| No Waveleting | 84.62 | NA | NA | NA | NA | NA | NA | NA | NA | NA | NA | NA |
| Wavelet (4.0 Hz) | 89.72 | 0.36 | 0.02 | 0.03 | 0.48 | 0.86 | 0.89 | 0.92 | 0.94 | 0.92 | 0.91 | 0.92 |
| Wavelet (2.0 Hz) | 88.33 | 0.44 | 0.02 | 0.45 | 0.84 | 0.86 | 0.89 | 0.92 | 0.94 | 0.92 | 0.91 | 0.92 |
| Wavelet (1.0 Hz) | 86.97 | 0.52 | 0.34 | 0.83 | 0.84 | 0.86 | 0.89 | 0.92 | 0.94 | 0.92 | 0.91 | 0.92 |
| Wavelet (0.5 Hz) | 87.20 | 0.59 | 0.74 | 0.83 | 0.84 | 0.86 | 0.89 | 0.92 | 0.94 | 0.92 | 0.91 | 0.92 |
| Wavelet (0.25 Hz) | 87.00 | 0.68 | 0.81 | 0.83 | 0.84 | 0.86 | 0.89 | 0.92 | 0.94 | 0.93 | 0.91 | 0.92 |
| Wavelet (0.10 Hz) | 86.54 | 0.75 | 0.82 | 0.83 | 0.84 | 0.86 | 0.89 | 0.92 | 0.94 | 0.92 | 0.91 | 0.92 |
| 4-month data |  |  |  |  |  |  |  |  |  |  |  |  |
| Hand-Edit | 69.55 | NA | NA | NA | NA | NA | NA | NA | NA | NA | NA | NA |
| No Waveleting | 74.02 | NA | NA | NA | NA | NA | NA | NA | NA | NA | NA | NA |
| Wavelet (4.0 Hz) | 86.78 | 0.21 | 0.02 | 0.02 | 0.27 | 0.85 | 0.83 | 0.80 | 0.84 | 0.86 | 0.86 | 0.84 |
| Wavelet (2.0 Hz) | 84.35 | 0.32 | 0.05 | 0.13 | 0.71 | 0.85 | 0.83 | 0.80 | 0.84 | 0.86 | 0.86 | 0.84 |
| Wavelet (1.0 Hz) | 81.70 | 0.49 | 0.19 | 0.35 | 0.82 | 0.85 | 0.84 | 0.81 | 0.85 | 0.87 | 0.87 | 0.85 |
| Wavelet (0.5 Hz) | 81.24 | 0.70 | 0.51 | 0.61 | 0.84 | 0.85 | 0.85 | 0.82 | 0.86 | 0.88 | 0.88 | 0.87 |
| Wavelet (.25 Hz) | 80.99 | 0.79 | 0.70 | 0.73 | 0.84 | 0.85 | 0.85 | 0.83 | 0.87 | 0.88 | 0.88 | 0.87 |
| Wavelet (.10 Hz) | 80.53 | 0.80 | 0.71 | 0.74 | 0.84 | 0.85 | 0.85 | 0.83 | 0.87 | 0.88 | 0.88 | 0.87 |
| 10-month data |  |  |  |  |  |  |  |  |  |  |  |  |
| Hand-Edit | 68.55 | NA | NA | NA | NA | NA | NA | NA | NA | NA | NA | NA |
| No Waveleting | 76.53 | NA | NA | NA | NA | NA | NA | NA | NA | NA | NA | NA |
| Wavelet (4.0 Hz) | 81.95 | 0.24 | 0.02 | 0.03 | 0.28 | 0.86 | 0.85 | 0.79 | 0.84 | 0.86 | 0.85 | 0.81 |
| Wavelet (2.0 Hz) | 81.24 | 0.33 | 0.05 | 0.13 | 0.70 | 0.86 | 0.85 | 0.79 | 0.84 | 0.86 | 0.85 | 0.81 |
| Wavelet (1.0 Hz) | 78.56 | 0.46 | 0.18 | 0.33 | 0.79 | 0.87 | 0.85 | 0.80 | 0.85 | 0.87 | 0.85 | 0.81 |
| Wavelet (0.5 Hz) | 81.62 | 0.61 | 0.44 | 0.54 | 0.80 | 0.87 | 0.86 | 0.80 | 0.85 | 0.87 | 0.85 | 0.82 |
| Wavelet (.25 Hz) | 81.56 | 0.68 | 0.58 | 0.63 | 0.80 | 0.87 | 0.86 | 0.80 | 0.85 | 0.87 | 0.85 | 0.82 |
| Wavelet (.10 Hz) | 81.62 | 0.69 | 0.59 | 0.64 | 0.81 | 0.87 | 0.86 | 0.80 | 0.85 | 0.87 | 0.85 | 0.82 |

**Table 4.** Statistics for the performance of various ICA rejection methods

| Method | Percent Files Retained | Average Percent Segments Retained | Average Percent ICs Rejected | Average Percent Variance Retained | r pre/post-Method |  |  |  |  |  |  |  |  |
| --- | --- | --- | --- | --- | --- | --- | --- | --- | --- | --- | --- | --- | --- |
|  |  |  |  |  | all data | 0.5 Hz | 1 Hz | 2 Hz | 5 Hz | 8 Hz | 12 Hz | 20 Hz | 30 Hz |
| Wavelet + ICA using MARA 0.5 | 30.43 | 31.56 | 98.26 | 1.62 | 0.19 | 0.05 | 0.05 | 0.08 | 0.10 | 0.10 | 0.11 | 0.08 | 0.10 |
| Wavelet + ICA using MARA 0.8 | 78.26 | 73.65 | 93.27 | 8.51 | 0.25 | 0.09 | 0.09 | 0.12 | 0.14 | 0.14 | 0.13 | 0.11 | 0.11 |
| Wavelet + ICA using iMARA 0.5 | 100.00 | 85.98 | 69.84 | 11.72 | 0.45 | 0.25 | 0.27 | 0.30 | 0.37 | 0.35 | 0.30 | 0.27 | 0.24 |
| Wavelet + ICA using iMARA 0.2 | 100.00 | 79.56 | 37.94 | 62.58 | 0.66 | 0.51 | 0.53 | 0.56 | 0.62 | 0.59 | 0.54 | 0.50 | 0.46 |
| Wavelet + ICA using ICLabel 0.5 | 100.00 | 85.02 | 76.72 | 29.78 | 0.46 | 0.25 | 0.26 | 0.28 | 0.32 | 0.26 | 0.24 | 0.19 | 0.17 |
| Wavelet + ICA using ICLabel 0.8 | 100.00 | 74.65 | 33.77 | 71.02 | 0.72 | 0.59 | 0.60 | 0.61 | 0.63 | 0.59 | 0.55 | 0.50 | 0.47 |

To test the efficacy of different bad channel detection functions and determine the optimal criterion values for detection, we compared a series of automated options to a set of expert-identified bad channels for twenty files from an EGI dataset (each file had the same subset of 39 channels evaluated). Specifically, the files were run through the HAPPE 1.0 legacy detection method for bad channels as well as several iterations of the Clean Rawdata function and combinations of Clean Rawdata with spectrum evaluation to optimize channel classification (shown in Table 2). Note that for iterations of Clean Rawdata with Flatline Criterion included, the Flatline default of 5 seconds was determined to be sufficient for detecting flat channels and was not manipulated further. We evaluated the outputs from each criterion for bad channel detection relative to the manually selected channels by summing the number of false negatives and false positives for each file and calculating the overall accuracy rate across files for that set of automated parameters. False negatives refer to channels that were manually marked as bad but not flagged as bad by the pipeline. False positives refer to channels that were manually marked as good but were marked bad by the pipeline. An extra emphasis was placed on finding the settings with high accuracy that produced the lowest number of false positives to avoid removing usable channels in the dataset. HAPPE+ER’s optimal settings produced 10 false negative channels and 9 false positive channels out of 780 channels across all 20 files for an overall accuracy rate of 97.6%.

### Artifact Rejection in Continuous Data

Many automated preprocessing pipelines include strategies to remove artifact from continuous EEG and ERP data prior to segmentation to reduce the number of artifact-laden trials that must be removed. Two of the most common such approaches include independent component analysis (ICA) and wavelet-thresholding. Briefly, ICA clusters data across electrodes into independent components that can segregate artifact from neural timeseries, while wavelet-thresholding parses data within frequency ranges using coefficients that can detect artifact fluctuations in either electrode data or independent components. For example, HAPPE 1.0 implements both a wavelet-enhanced ICA (W-ICA) approach where wavelet thresholding is used to remove artifact from independent components, followed by a second ICA decomposition where artifact component timeseries are identified and removed. To optimize artifact detection and removal for ERP analyses, we evaluated a number of contemporary approaches using ICA and wavelet-thresholding, namely: wavelet-enhanced ICA (W-ICA), wavelet-thresholding without ICA, W-ICA followed by ICA (as in HAPPE 1.0), ICA, and for comparison, an approach without continuous data artifact rejection steps prior to ERP segmentation (as in traditional ERP-preprocessing) with automated segment rejection, and an approach with no continuous data artifact rejection prior to segmentation with manual segment rejection, the current gold standard ERP processing approach.

To evaluate the performance of each method in rejecting artifacts from continuous data, we examined both adult ERP data (a dataset with very clean signals), as well as developmental ERP data at both 4- and 10-months of age (generally with higher levels of artifact contamination) from both healthy babies and those with complex medical histories including early general anesthesia experiences. Parental consent for infants and consent for adults was obtained in accordance with Boston Children’s Hospital’s IRB. Specifically, we evaluated a highly stereotyped and robust ERP across datasets, the visual evoked potential (Sokol, 1976; Varcin and Nelson, 2016). The adults (n = 24) underwent a grating-VEP paradigm while the infants at both ages underwent a pattern-reversal VEP paradigm (n = 23 at 4-months, n = 17 at 10-months). Each dataset was run through HAPPE+ER in batch for all continuous data rejection approaches described above.

The artifact-rejection approaches were carried out as follows. W-ICA includes first performing an ICA decomposition of the EEG signal into components, after which all components’ timeseries are subjected to wavelet transform and thresholded to remove artifact before all the components’ cleaned timeseries are translated back to EEG channel data (Castellanos and Makarov, 2006). W-ICA followed by ICA includes a second ICA decomposition with automated component rejection (via MARA in adult data and iMARA in the developmental data). Wavelet-thresholding may also be employed independent of ICA, such that each channel’s timeseries undergoes thresholding for artifact instead of independent components from the ICA decomposition. Wavelet-thresholding with ICA includes initial wavelet thresholding followed by ICA with automated component rejection. ICA with automated component rejection may also be employed without waveleting. Finally, we examined removing all instances of wavelet-thresholding and ICA entirely (i.e., no artifact removal in continuous data) to match traditional preprocessing for ERP analyses. The data was then subjected to either automated segment rejection (as for the prior approaches) or to manual segment rejection, carried out by an expert with over a decade of experience in manually editing EEG and ERP data.

There are several important parameters that require further consideration when using both ICA and waveleting with ERP analyses. First, note that ICA typically requires a 1 Hz high-pass filter for adequate ICA performance (Winkler et al., 2015), which introduces a barrier to ERP analyses that include data below 1 Hz. To evaluate ICA within the context of ERP analyses, we took the following steps. First, each unfiltered file was copied, and the copy was subjected to a 1 Hz high-pass filter. Processing proceeded on the data copy for the approaches using W-ICA and/or ICA. As appropriate, the wavelet-thresholded artifact timeseries and/or the rejected artifact-laden independent component timeseries were then subtracted from the unfiltered file, effectively removing all waveleting- and/or ICA-identified artifacts from the data. The unfiltered file was then carried forward in analysis as the cleaned data eligible for further processing steps (e.g., ERP filtering, segmentation).

The second consideration for both ICA and waveleting approaches is that both required initial optimization for the ERP datasets. For waveleting, the optimum wavelet thresholding resolution for ERP analyses in both adult and developmental populations needed to be determined. Similarly, for ICA, the optimum algorithm for automated IC rejection with infant data was unclear, especially as multiple options have recently been released (e.g. Adjusted Adjust (Leach et al., 2020), iMARA (Haresign et al., 2021)). Therefore, both wavelet resolution and IC rejection steps were first optimized for subsequent method comparisons as described below. All statistical comparisons in subsequent sections were conducted as repeated-measures ANOVAs with post-hoc pairwise comparisons in SPSS software version 27.

#### Optimization of wavelet thresholding resolution for ERP analysis

Wavelet thresholding decomposes the data out into frequency ranges dictated by a resolution parameter, but the appropriate resolution level for ERP analyses has not been validated before. The resolution level is critical as it determines the bins of frequencies that are considered together for identifying and removing artifacts. Therefore, across adult and developmental datasets, we examined how wavelet resolution level influenced the VEP waveform, number of segments retained, and the degree of change in the data across frequencies (via quality control cross-correlation values) to optimize this approach for ERP analyses before testing approaches involving wavelet-thresholding. Specifically, we evaluated decomposing the data so that the finest frequency resolution was one of the following: ≤ ∼ 4 Hz, ≤ ∼ 2 Hz, ≤ ∼ 1 Hz, ≤ ∼ 0.5 Hz, ≤ ∼ 0.25 Hz, or ≤ ∼ 0.1 Hz.

Across adult and developmental datasets, the 4hz resolution level visibly shrunk the VEP waveform relative to the other parameters, suggesting worse fit for ERP data. In the 4-month data, the ≤ 2 Hz resolution level also resulted in altered VEP waveform morphology relative to the other wavelet levels and approaches (manual and automated segment rejection). The other resolution levels from ≤ 1 Hz to ≤ 0.1 Hz did not result in meaningful distortion of the ERP waveform relative to approaches without waveleting through initial visual inspection.

The effect of wavelet resolution level on trial retention rates was also considered across datasets. There were minimal differences between approaches for the adult VEP trial retention rates (F(7) = 2.18, p = 0.039, ƞ_*p*_^2^ = 0.086). No wavelet resolution level resulted in significantly different trial retention rates from the manually edited rate (all p > 0.05). Wavelet resolution levels from ≤ 1 Hz down to ≤ 0.1 Hz were not significantly different from each other in trial retention rates (all p > 0.05). This pattern of results suggests that waveleting approaches at most resolutions considered for optimization perform comparably each other and to the gold-standard manual editing approach for trial retention for low-artifact data. There were similarly minimal differences between resolution levels for both 4-month and 10-month developmental VEP trial retention rates (4-month: F(7) = 13.265, p = 2.5*10^-13^, ƞ_*p*_^2^ = 0.376; 10-month: F(7) = 5.736, p = 1.1*10^-5^, ƞ_*p*_^2^ = 0.112). As in the adult data, wavelet resolutions of ≤ 0.1 Hz, 0.25 Hz, and 0.5 Hz were not significantly different from each other in the number of retained trials in either developmental dataset (all p > 0.05). However, in both developmental datasets, manual editing resulted in significantly fewer retained trials compared to wavelet thresholded data at almost every resolution level (p < 0.05). This pattern of results suggests that waveleting across resolution levels does not impact trial retention meaningfully but outperforms the gold-standard manual editing approach for rates of trial retention in higher-artifact developmental data.

Finally, the degree of signal change during the wavelet-thresholding process was compared across wavelet resolution levels using the cross-correlation values for the EEG signal before and after wavelet-thresholding. There were striking and consistent effects of wavelet resolution level on the correlation of the EEG signal pre- and post-waveleting across adult and developmental datasets (adult: F(5) = 214.983, p = 3.98*10^-58^, ƞ_*p*_^2^ = 0.9; 4-month: F(5) = 822.939, p = 2.20*10^-85^, ƞ_*p*_^2^ = 0.743; 10-month: F(5) = 193.496, p = 3.69*10^-43^, ƞ_*p*_^2^ = 0.924). In each dataset, the more fine-grained the resolution, the less the EEG signal changed during wavelet-thresholding (significantly higher cross-correlation values were observed). Examining the cross-correlations at specific key frequencies in the signal confirmed that this effect was driven by change specifically in the low-frequencies as expected, since changing the wavelet resolution within the range we tested impacts treatment of the low frequency data. Notably, the finest resolution level of ≤ 0.1 Hz produced signal change the most comparable across frequencies, especially in the clean adult data with less low-frequency signal drift.

Given this pattern of results across waveform inspection, trial retention, and signal change during processing across datasets, the wavelet resolution level was set to ≤ 0.1 Hz to optimize wavelet-thresholding performance in ERP analyses.

#### Optimization of Independent Component Automated Rejection for Developmental ERP analyses

For methods that use ICA, multiple algorithms exist that perform automated component rejection, but few have been tested or validated in developmental ERP data (though see iMARA (Haresign et al., 2021), Adjusted ADJUST (Leach et al., 2020), and ICLabel (Pion-Tonachini et al., 2017)). Therefore, we first compared across algorithms and rejection thresholds to optimize settings for ERP analyses in infant data using the 4-month VEP dataset. We selected the 4-month dataset for infant optimization instead of the 10-month dataset as the VEP is actively developing across all waveform components at 4 months of age and is more distinct from adult VEP morphology where prior validation exists. We compared the following automated rejection algorithms: Adjusted ADJUST, ICLabel, MARA, and iMARA. Adjusted ADJUST has previously been validated on infant data using an EGI high-density system as used for collecting the present datasets, so the default threshold settings from that validation were used here. For ICLabel, MARA, and iMARA algorithms without prior validation on the same system as this dataset, they were each evaluated at the standard rejection threshold of 0.5 or higher probability of being an artifact component and at a liberal threshold for data retention that only rejected components with 0.8 or higher probability of being artifact. Both the MARA (at both 0.5 and 0.8 thresholds) and the Adjusted ADJUST algorithms rejected multiple files (known to have usable VEP data as determined by manual inspection by an experienced researcher), so these algorithms were not considered further in primary analyses. Secondary analyses including Adjusted ADJUST on the subset of participants with retained data are referenced in brief and provided in the supplemental data tables. For primary analyses, the ICLabel 0.5, ICLabel 0.8, iMARA 0.5, and iMARA 0.8 algorithm-threshold combinations were compared statistically with manual editing and the segment rejection only approach in their effects on 1) participant and trial retention rates, 2) percent data retained post-processing, and 3) VEP morphology.

First, with respect to participant and trial retention, while all four algorithm-threshold combinations preserved 100% of the participants, there were significant differences in trial retention rates between these methods (F(5) = 13.491, p = 2.93*10^-10^, ƞ_*p*_^2^ = 0.380). All automated approaches retained more trials than the manual editing option (all p < 0.05), and ‘automated segment rejection only’ retained significantly fewer trials than iMARA 0.5, iMARA 0.2, and ICLabel 0.5 options (but was no different from ICLabel 0.8 in retention rates). The iMARA 0.5 and ICLabel 0.5 combinations retained significantly more trials than all other options (all p < 0.05) (but were not significantly different from each other, p > 0.05). Secondary analyses on the subset of participants retained by Adjusted ADJUST revealed that this algorithm retained significantly fewer trials than iMARA 0.2, iMARA 0.5, ICLabel 0.5, and ‘automated segment rejection only’ (all p < 0.05) and was not significantly different from trial retention rates of manual editing or ICLabel 0.8 (all p > 0.05). Thus, for trial retention, iMARA and ICLabel approaches both performed equally best with standard artifact thresholds (rather than liberal thresholds).

Second, the algorithm-threshold combinations were statistically compared on the percent data variance retained post-processing. This comparison did not include the manual editing or segment rejection only approaches since they do not remove variance from the continuous data prior to trial rejection. There was a statistically significant effect of algorithm-threshold combination on how much data was retained in the course of preprocessing (F(3) = 90.263, p = 2.588*10^-23^, ƞ_*p*_^2^ = 0.804). As expected, we found that both iMARA and ICLabel algorithms with liberal thresholds retained significantly more data variance post-processing than the algorithms with standard thresholds (all p < 0.05). Moreover, ICLabel 0.5 retained significantly more variance than the iMARA 0.5 option (p < 0.05). Thus, the liberal rejection thresholds did result in increased data retention during processing.

Finally, the effects of algorithm-threshold combinations were evaluated with respect to VEP peak amplitude morphology. Notably, the number of trials within an individual dataset used to generate the average VEP waveform affects the peak waveform amplitude (Luck, 2014), so for the following visualizations and statistical comparisons of VEP morphology, wherever automated methods had preserved greater numbers of trials relative to manual editing, a random subset of trials from each individual were selected to match the number of trials from manual editing for that same individual. There was a corresponding, consistent statistical pattern of algorithm-threshold effects on the VEP morphology across the N1, N1-P1 peak-to-peak, and P1-N2 peak-to-peak component amplitudes (N1: F(5) = 15.235, p = 2.33*10^-11^, ƞ_*p*_^2^ = 0.409; N1-P1: F(5) = 19.805, p = 4.94*10^-14^, ƞ_*p*_^2^ = 0.474; P1-N2: F(5) = 14.586, p = 5.9*10^-11^, ƞ_*p*_^2^ = 0.399). For each component amplitude, the manual rejection and automated segment rejection approaches were no different from each other but significantly larger than amplitudes from all MARA, iMARA, and ICLabel options (all p < 0.05). The iMARA and ICLabel approaches with the liberal 0.8 thresholds were no different from each other but were significantly larger than iMARA and ICLabel approaches with standard 0.5 thresholds (all p < 0.05). That is, while all component rejection approaches reduced ERP amplitudes relative to the gold-standard manual editing approach, the algorithms with liberal thresholds introduced less amplitude shrinkage (via retaining a greater percent of the data) than algorithms with standard thresholds for rejection.

This pattern of results suggests that iMARA and ICLabel achieve comparable performance across measures in developmental ERP analyses. However, automated component rejection performance, regardless of algorithm, depends highly on the rejection threshold selected. That is, there appears to be a tradeoff such that the standard rejection thresholds enable increased trial retention rates but remove a larger percent of the data and so result in more shrunken ERP amplitudes than the liberal rejection thresholds. Without a clear optimal algorithm-threshold option, we proceeded with the iMARA 0.2 combination for automated component rejection in the developmental datasets as this algorithm was one of the two better-performing options and was generated specifically for developmental ICA analyses.

### Comparison of artifact rejection approaches for ERP analyses

Following optimizations, the following approaches were compared for artifact removal to determine the strategy best suited for ERP analyses across the lifespan: manual editing, automated trial rejection without prior artifact removal in the continuous signal, ICA, wavelet thresholding, wavelet thresholding with ICA, wICA, and wICA with ICA. Methods were evaluated on how they impacted the VEP waveform morphology, rate of participant and trial retention, and degree of signal change and data retention during processing. As before, for visualizations and statistical comparisons of VEP morphology, wherever automated methods had preserved greater numbers of trials relative to manual editing, a random subset of trials from each individual were selected to match the number of trials from manual editing for that same individual. Methods are compared below first for low-artifact adult VEP data and then in two higher-artifact developmental, clinical VEP datasets.

#### Adult ERP Comparisons

The different artifact rejection methods were first compared across participant and trial retention rates in the adult VEP data. For all ICA-based approaches in the adult data, MARA with default settings was used for component rejection (components with artifact probability greater than 0.5 were removed). All methods retained 100% of the participants in the adult sample (data remained post-processing for all individuals). That is, there was no erroneous sample attrition regardless of processing strategy. A statistically significant effect of artifact rejection method on trial retention rate was observed (F(6) = 2.914, p = 0.010, ƞ_*p*_^2^ = 0.112). Post-hoc comparisons revealed that no approach was significantly different from the manual editing trial retention rate (all p > 0.05). Within the set of automated rejection approaches, wICA resulted in significantly fewer trials retained compared to all other approaches (all p < 0.05), and wICA with ICA resulted in fewer trials than ICA only (p = 0.003). This pattern of results suggests that in low-artifact, adult data, most automated artifact rejection approaches (aside from wICA) are comparable to each other and to manual editing in terms of trial retention.

Next, the degree of signal change and data retention rate during processing were compared across methods in this very clean dataset. Notably, the ICA-based methods all retained less than 50% of the data post-processing (Table 5). Correspondingly, the ICA-based methods demonstrated significantly lower correlations between the pre- and post-processed data compared to waveleting-based approaches across frequencies (F(4) = 66.625, p = 2.37*10^-26^, ƞ_*p*_^2^ = 0.743). There were no significant differences within the ICA-based methods (ICA, waveleting with ICA, wICA with ICA) in the correlations pre- and post-processing (all p > 0.05). There were no significant differences within the waveleting approaches (wavelet thresholding, wICA) in the correlations pre- and post-processing either (p > 0.05). This pattern of results in the clean adult data suggest the wavelet-based methods preserved substantially more of the underlying signal during processing compared with ICA-based approaches.

**Table 5.** Statistics for the performance of various artifact rejection methods on adult data.

| Method | Percent Files Retained | Average Percent Segments Retained | Average Percent ICs Rejected | Average Percent Variance Retained | r pre/post-Method |  |  |  |  |  |  |  |  |
| --- | --- | --- | --- | --- | --- | --- | --- | --- | --- | --- | --- | --- | --- |
|  |  |  |  |  | all data | 0.5 Hz | 1 Hz | 2 Hz | 5 Hz | 8 Hz | 12 Hz | 20 Hz | 30 Hz |
| Hand-Edit | 100.00 | 86.08 | NA | NA | NA | NA | NA | NA | NA | NA | NA | NA | NA |
| No Waveleting, No ICA | 100.00 | 84.62 | NA | NA | NA | NA | NA | NA | NA | NA | NA | NA | NA |
| ICA Only | 100.00 | 95.24 | 63.75 | 22.16 | 0.45 | 0.30 | 0.35 | 0.38 | 0.47 | 0.58 | 0.64 | 0.56 | 0.46 |
| Waveleting Only | 100.00 | 84.92 | NA | NA | 0.75 | 0.82 | 0.83 | 0.84 | 0.86 | 0.89 | 0.92 | 0.94 | 0.92 |
| Waveleting and ICA | 100.00 | 92.06 | 60.11 | 40.62 | 0.47 | 0.34 | 0.35 | 0.40 | 0.51 | 0.60 | 0.65 | 0.59 | 0.48 |
| wICA Only | 100.00 | 84.13 | NA | NA | 0.78 | 0.71 | 0.71 | 0.66 | 0.71 | 0.82 | 0.88 | 0.91 | 0.89 |
| wICA and ICA | 100.00 | 86.24 | 54.58 | 47.07 | 0.46 | 0.34 | 0.36 | 0.38 | 0.50 | 0.63 | 0.69 | 0.63 | 0.53 |

**Table 6.** Statistics for the performance of various artifact rejection methods on 4-month and 10-month data.

| Method | Percent Files Retained | Average Percent Segments Retained | Average Percent ICs Rejected | Average Percent Variance Retained | r pre/post-Method |  |  |  |  |  |  |  |  |
| --- | --- | --- | --- | --- | --- | --- | --- | --- | --- | --- | --- | --- | --- |
|  |  |  |  |  | all data | 0.5 Hz | 1 Hz | 2 Hz | 5 Hz | 8 Hz | 12 Hz | 20 Hz | 30 Hz |
| 4-Month Data |  |  |  |  |  |  |  |  |  |  |  |  |  |
| Hand-Edit | 100.00 | 69.55 | NA | NA | NA | NA | NA | NA | NA | NA | NA | NA | NA |
| No Waveleting, No ICA | 100.00 | 74.02 | NA | NA | NA | NA | NA | NA | NA | NA | NA | NA | NA |
| ICA Only | 100.00 | 77.13 | 46.60 | 45.74 | 0.66 | 0.52 | 0.52 | 0.53 | 0.60 | 0.58 | 0.55 | 0.48 | 0.43 |
| Waveleting Only | 100.00 | 80.53 | NA | NA | 0.80 | 0.71 | 0.74 | 0.84 | 0.86 | 0.84 | 0.81 | 0.85 | 0.87 |
| Waveleting and ICA | 100.00 | 79.56 | 37.94 | 62.58 | 0.66 | 0.51 | 0.53 | 0.56 | 0.62 | 0.59 | 0.54 | 0.50 | 0.46 |
| wICA Only | 100.00 | 60.44 | NA | NA | 0.82 | 0.73 | 0.74 | 0.76 | 0.82 | 0.82 | 0.82 | 0.87 | 0.89 |
| wICA and ICA | 100.00 | 74.61 | 19.71 | 77.46 | 0.71 | 0.57 | 0.58 | 0.61 | 0.69 | 0.69 | 0.66 | 0.65 | 0.64 |
| 10-Month Data |  |  |  |  |  |  |  |  |  |  |  |  |  |
| Hand-Edit | 100.00 | 69.97 | NA | NA | NA | NA | NA | NA | NA | NA | NA | NA | NA |
| No Waveleting, No ICA | 100.00 | 76.53 | NA | NA | NA | NA | NA | NA | NA | NA | NA | NA | NA |
| ICA Only | 100.00 | 72.16 | 20.59 | 73.70 | 0.85 | 0.78 | 0.78 | 0.79 | 0.84 | 0.87 | 0.79 | 0.72 | 0.70 |
| Waveleting Only | 100.00 | 81.62 | NA | NA | 0.69 | 0.59 | 0.64 | 0.81 | 0.87 | 0.86 | 0.80 | 0.85 | 0.87 |
| Waveleting and ICA | 100.00 | 73.03 | 6.79 | 92.75 | 0.78 | 0.71 | 0.73 | 0.76 | 0.84 | 0.83 | 0.76 | 0.78 | 0.77 |
| wICA Only | 100.00 | 71.66 | NA | NA | 0.76 | 0.67 | 0.67 | 0.71 | 0.83 | 0.82 | 0.76 | 0.83 | 0.83 |
| wICA and ICA | 100.00 | 73.47 | 4.45 | 94.30 | 0.73 | 0.63 | 0.65 | 0.69 | 0.81 | 0.81 | 0.74 | 0.78 | 0.78 |

Finally, artifact rejection methods were evaluated on their impact on the adult VEP morphology using trial number matched datasets. In the context of this pristine dataset, all the automated artifact rejection approaches tested preserved gross morphology as determined by visual inspection (Figure 4). However, repeated measures ANOVAs comparing the three VEP components’ peak values (N1, N1-P1 peak-to-peak, and P1-N2 peak-to-peak amplitudes) revealed a pattern of statistical differences between rejection methods (N1: F(6) = 2.434, p = 0.029, ƞ_*p*_^2^ = 0.096; N1-P1: F(6) = 7.003, p = 2*10^-6^, ƞ_*p*_^2^ = 0.233; P1-N2: F(6) = 10.801, p = 7.83*10^-10^, ƞ_*p*_^2^ = 0.320). Specifically, the wavelet thresholding method generated a more negative N1 component amplitude than most other methods (except wICA and segment rejection only methods, p > 0.05). No other methods produced N1 amplitudes that were significantly different from each other (all p > 0.05). Across the N1-P1 and P1-N2 components, wavelet-based approaches (wavelet and wICA) resulted in amplitudes that were not statistically different from amplitudes generated by the manual editing and segment rejection only methods (all p > 0.05), while all ICA-based approaches (ICA, waveleting with ICA, wICA with ICA) resulted in significantly smaller peak-peak amplitudes (all p < 0.05). Within the ICA-based approaches, ‘wICA with ICA’ and ICA resulted in amplitudes that were significantly larger than ‘wavelet with ICA’ amplitudes. This pattern of results in the clean adult data suggests that the ICA-based approaches employed here reduced ERP amplitudes, while the wavelet-based approaches tested here did not shrink ERP amplitudes.

Together this pattern of results in the low-artifact adult VEP data demonstrates that of the automated artifact rejection approaches, the wavelet thresholding method best-preserved VEP morphology and the EEG signal during processing with trial retention rates no different from either gold-standard manual editing or other automated options. Notably, this adult dataset was intentionally particularly pristine (experienced EEG researchers were included in the sample and the paradigm was brief). Thus, while there was no observed benefit here to wavelet-thresholding relative to the ‘segment rejection only’ approach (no artifact rejection in continuous data prior to trial rejection), including artifact rejection in continuous data should provide benefits in adult ERP data more broadly, and these data suggest wavelet-thresholding provides the best artifact rejection approach of those tested.

The different artifact rejection methods were also evaluated on their impact on two developmental VEP datasets from 4- and 10-month-olds that included more artifacts during recording than the healthy adult dataset. For all ICA-based approaches in the developmental data, the infant-trained iMARA algorithm with the liberal data-retention setting was used for component rejection (components with artifact probability greater than 0.8 were removed). As before, methods were evaluated on their effects on participant and trial retention and the VEP morphology.

All methods tested on both 4- and 10-month datasets retained 100 percent of the participants. That is, there was no erroneous sample attrition regardless of processing strategy. A statistically significant effect of artifact rejection method on trial retention rate was observed in both the 4- and 10-month datasets, however (4-month: F(6) = 15.585, p = 1.82*10^-13^, ƞ_*p*_^2^ = 0.415; 10-month: F(6) = 6.557, p = 8*10^-6^; ƞ_*p*_^2^ = 0.291). Specifically, in the 4-month data, manual editing retained significantly fewer trials than all other methods except wICA (where manual editing retained significantly more trials, p < 0.05). Wavelet thresholding preserved significantly more trials than all other methods except waveleting with ICA (no difference, p > 0.05).

Automated segment rejection (without any prior artifact removal) preserved significantly more trials than wICA but was no different from the trial retention rate of wICA with ICA (p > 0.05), and retained significantly fewer trials than waveleting, ICA, and waveleting with ICA. That is, in the 4-month data, there was a trial retention benefit to using some methods of continuous artifact rejection prior to segment rejection, and in particular wavelet thresholding preserved the most trials of all the methods tested. Similarly, in the 10-month data, wavelet thresholding preserved more trials than all other methods tested (all p < 0.05). Automated segment rejection (without prior artifact removal) preserved the second-highest number of trials, significantly more than all other methods except wavelet thresholding (all other p < 0.05). Manual editing trial retention rates were not statistically different from those using ICA, waveleting with ICA, wICA, or wICA with ICA (all p > 0.05). Thus, across both 4-month and 10-month datasets, wavelet thresholding consistently preserved more trials per participant than any other method.

Next, methods were compared with respect to effects on VEP morphology across 4- and 10-month datasets using trial number matched datasets (matched to manual editing trial retention values). In the 4-month dataset there was a consistent pattern of effects across VEP components (N1: F(6) = 5.426, p = 4.8*10^-5^, ƞ_*p*_^2^ = 0.198; N1-P1: F(6) = 12.786, p = 2.39*10^-11^, ƞ_*p*_^2^ = 0.368; P1-N2: F(6) = 8.410, p = 9.89*10^-8^, ƞ_*p*_^2^ = 0.277). Specifically, waveleting methods (wavelet thresholding and wICA) resulted in amplitudes that were not significantly different from those produced by either manual editing or automated segment rejection without prior artifact removal (p > 0.05; except for N1-P1 amplitudes after waveleting were significantly smaller than manual editing amplitudes but not those from automated segment rejection only), whereas ICA-based approaches resulted in smaller amplitudes compared to manual editing and automated segment rejection (p < 0.05). There was less differentiation between methods in the 10-month data sample. The omnibus F test was not significant for the test of differences in the N1 peak amplitude across methods (F(6) = 0.343, p = 0.913, ƞ_*p*_^2^ = 0.021). Similarly, for the N1-P1 peak-to-peak amplitudes (F(6) = 6.551, p = 0.029, ƞ_*p*_^2^ = 0.134), the amplitudes derived from manual editing and automated segment rejection were not significantly different from any other methods (all p > 0.05). P1-N2 peak-to-peak amplitudes were significantly different across methods (F(6) = 3.692, p = 0.002, ƞ_*p*_^2^ = 0.188). Namely, manual editing produced larger peak-to-peak amplitudes than automated segment rejection only and wavelet thresholding but not other methods (all p > 0.05). Automated approaches were largely not significantly different from each other though, except for wICA with ICA that produced a larger P1-N2 amplitude than other approaches besides manual editing and wavelet thresholding with ICA (p > 0.05).

Together this pattern of results in the developmental and adult VEP data demonstrates that wavelet-thresholding offers the best solution of the methods tested here for ERP processing across the lifespan. Wavelet-thresholding preserved more trials than other approaches across developmental datasets and in clean adult data retained trials at the same high rate as other approaches. Wavelet-thresholding also generally better preserved VEP amplitudes relative to the other automated artifact rejection approaches in both developmental and adult data, resulting in amplitudes consistent with the current field-standard from manual-editing. Moreover, wavelet-thresholding offers a computationally efficient and fast method of artifact rejection compared to the other automated approaches tested here. Thus, even in cases where there may be no distinct ERP-morphology or trial retention differences between wavelet-thresholding and ICA-based approaches, wavelet-thresholding still provides the most tractable solution from an efficiency viewpoint. HAPPE+ER therefore implements the wavelet-thresholding method tested here as its approach for artifact rejection in continuous data for ERP analyses across the lifespan.

### Segmentation (Optional)

HAPPE+ER includes an optional data segmentation step for ERP analyses in which data is segmented around the events using the stimulus onset tags specified by the user. HAPPE+ER inherently corrects for any timing offset as part of the segmentation process, using the offset input by the user at the start of the HAPPE+ER run (if no timing offset is present, user may specify a zero-millisecond offset during that process). Two additional artifact-reduction options are available for ERPs if segmentation is selected: baseline correction and data interpolation within segments, described in detail below. Users may segment their data regardless of whether these other options are enabled (see Figure 1 for complete diagram of optional segment-related options in HAPPE+ER).

**Figure 1.**
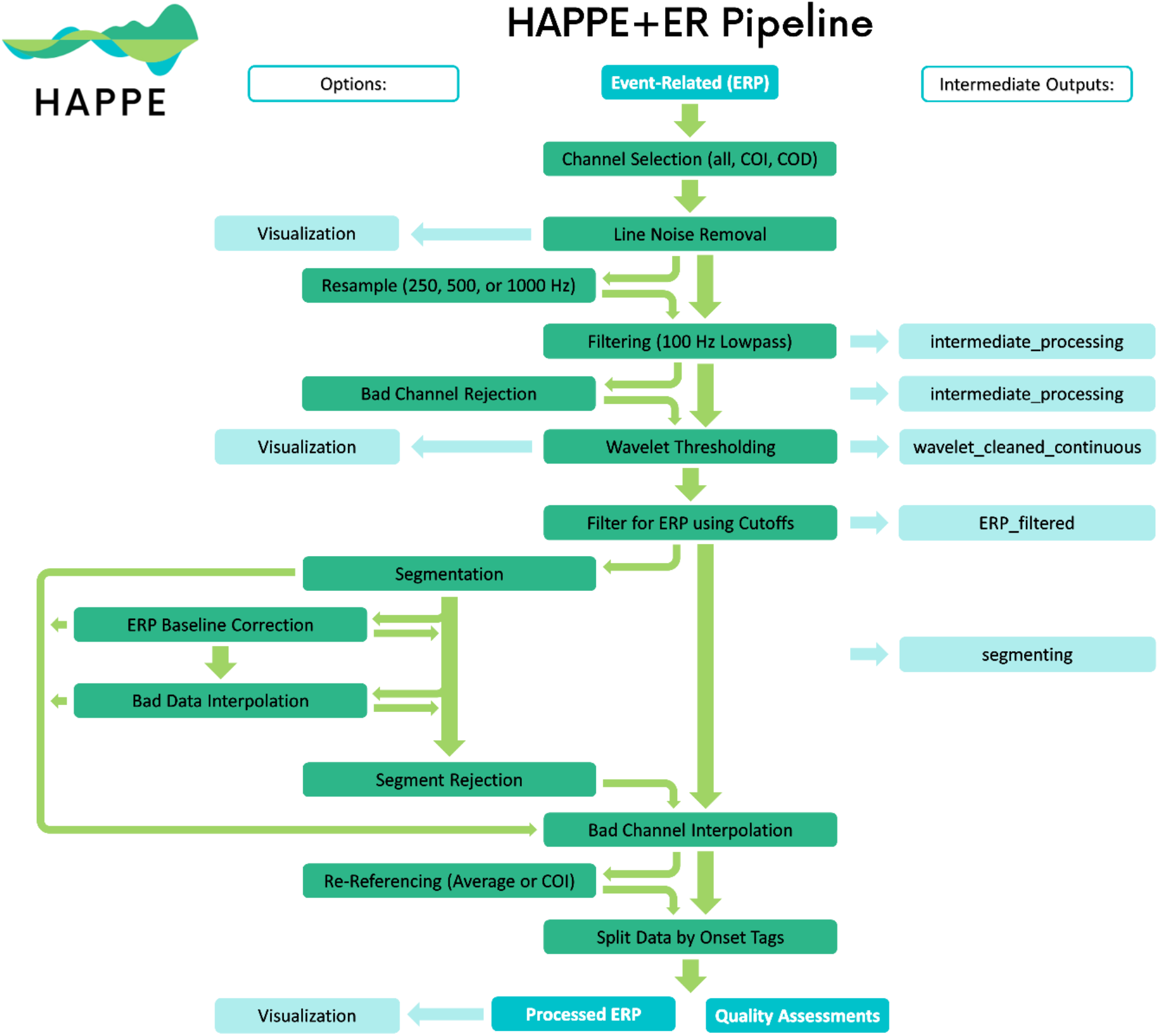
Image illustrating the HAPPE+ER pipeline’s processing steps. Intermediate outputs are noted by the light blue boxes on the right and are labeled according to the folder where they are saved. User options are displaced to the left, with bright green arrows indicating all possible methods of flow between options.

**Figure 2.**
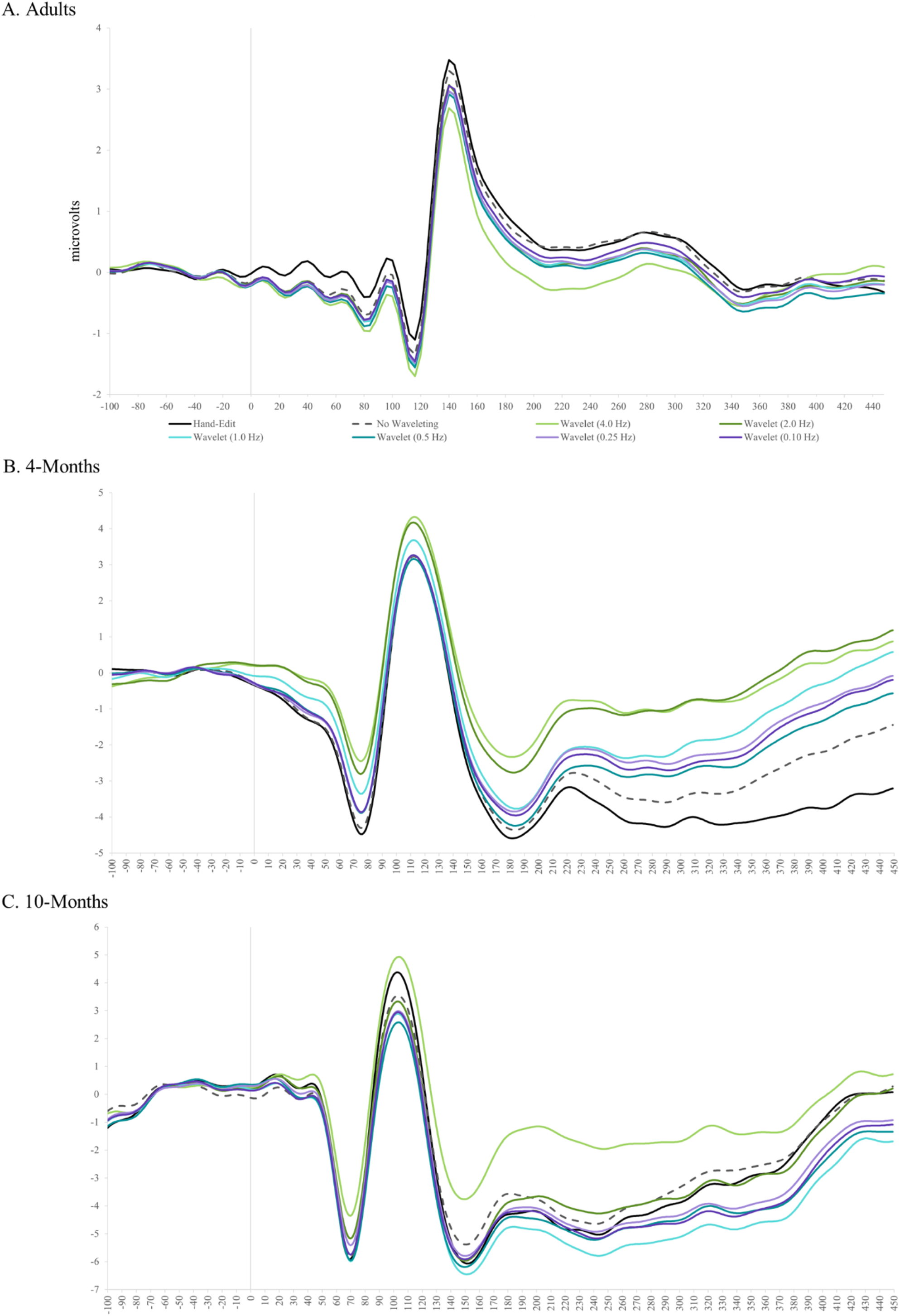
Three images illustrating the resultant VEP ERP waveform following processing using an array of wavelet levels, via hand-editing, and through the HAPPE+ER pipeline without artifact rejection steps, on adult (A), 4-month (B), and 10-month (C) data

**Figure 3.**
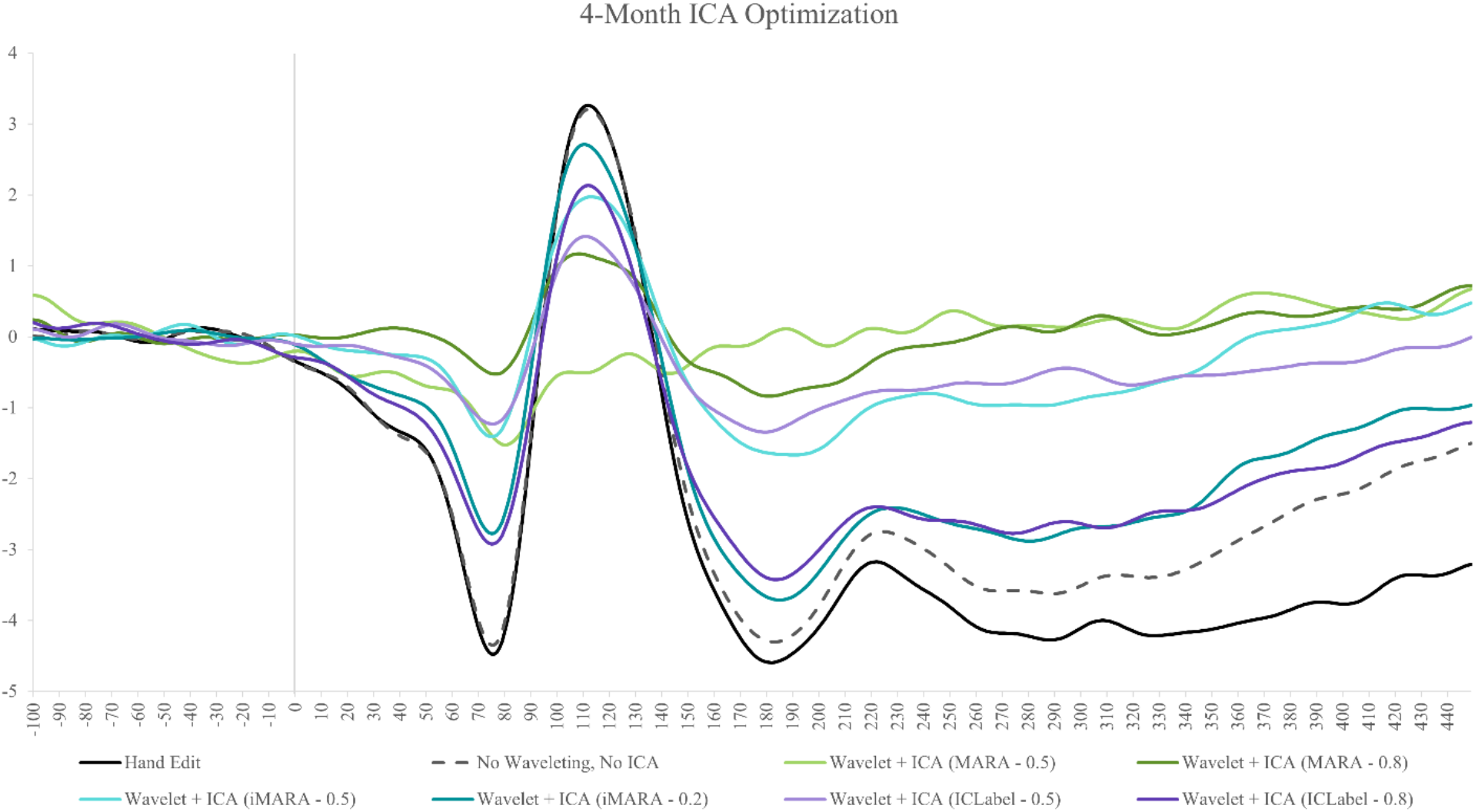
Figure illustrating the VEP ERP waveform generated after processing 4-month data using various ICA rejection methods, through hand-editing, and with segment rejection only. Greens indicate processing with MARA, blues with iMARA, and purples with ICLabel. Darker hues indicate a more liberal rejection threshold.

**Figure 4.**
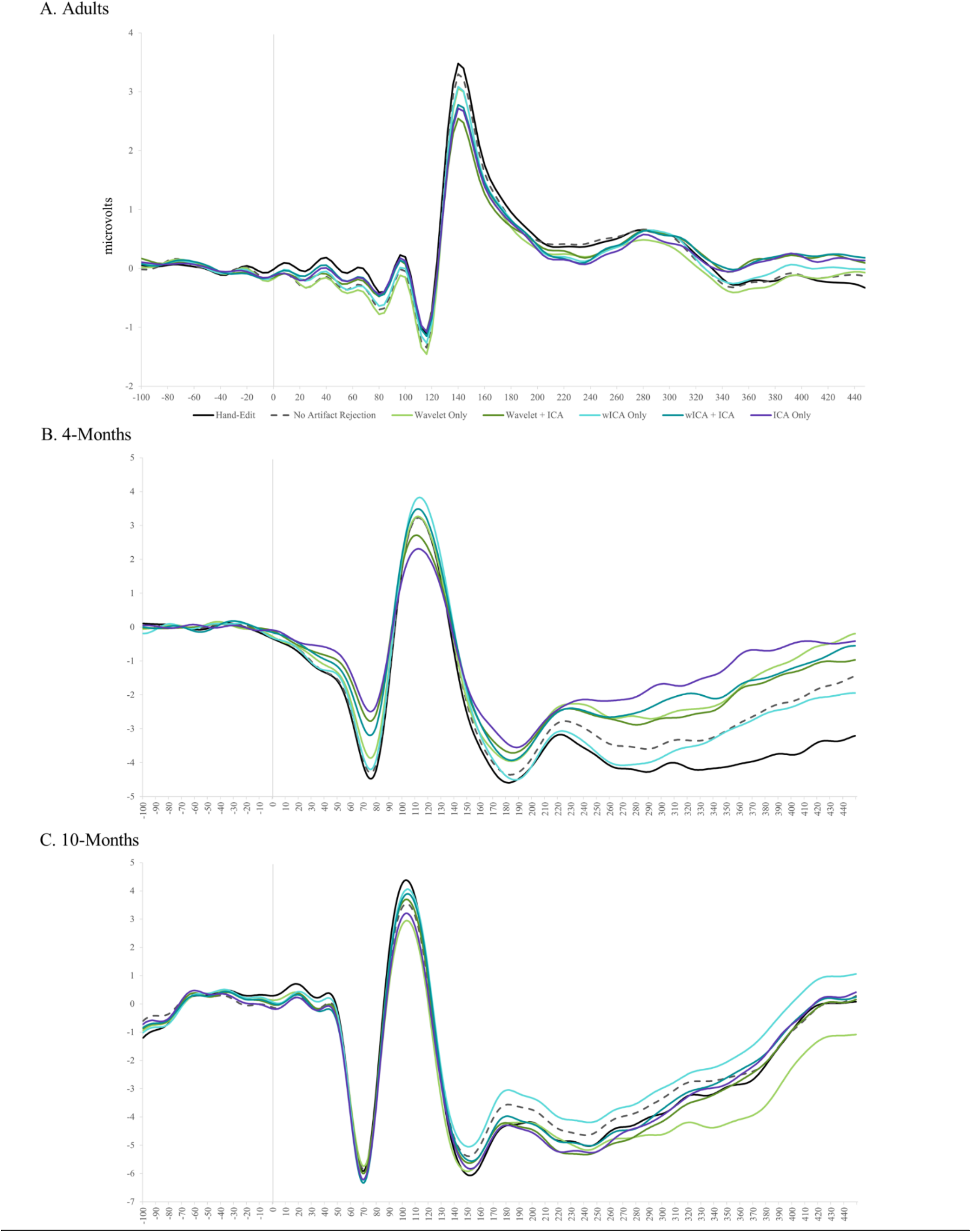
Three images illustrating the resultant VEP ERP waveform following processing using an array of artifact rejection methods and segment rejection only, on adult (A), 4-month (B), and 10-month (C) data.

### Baseline Correction (Optional)

Users may perform baseline correction on segmented ERP data to correct drift-related differences between data segments. HAPPE+ER uses a standard mean subtraction method to achieve this correction where the mean value from the user-specified baseline window (defined relative to the stimulus onset) is subtracted from the data in the post-stimulus window. This step is implemented via the rmbase function in EEGLAB. The baseline correction option may be especially useful for users with saline-based acquisition systems and/or longer recording periods where drift is more commonly observed.

### Data Interpolation within Segments (Optional)

For datasets where segment rejection would lead to an unacceptably low remaining number of segments for ERP analysis, users may choose an optional post-segmentation step involving the interpolation of data within individual segments for channels determined to be artifact-contaminated during that segment, as implemented by FASTER software (Nolan et al., 2010). Each channel in each segment is evaluated on the four FASTER criteria (variance, median gradient, amplitude range, and deviation from mean amplitude), and the Z score for each channel in that segment is generated for each of the four metrics. Any channels with one or more Z scores that are greater than 3 standard deviations from the mean for an individual segment are marked bad for only that segment. These criteria may identify segments with residual artifacts in specific channels. Subsequently, for each segment, the channels flagged as bad in that segment have their data interpolated with spherical splines, as in FASTER. This allows users to maintain the maximum number of available segments, while still maximizing artifact rejection within individual segments. However, we caution users from implementing this option in cases where they have selected channel subset such that the channels are distributed with significant distance between them as the interpolation process would pull data from distal channels that does not reflect the appropriate activity profile for that scalp space.

### Segment Rejection (Optional)

For pre-segmented data or data run through HAPPE+ER’s optional segmentation step, users can choose to reject segments. If selected, segments can be rejected based on amplitude, joint probability, or both criteria. Amplitude-based rejection is useful for removing residual high-amplitude artifacts (e.g., eye blinks, drift from drying electrodes, discontinuities). If selected, users specify an artifact amplitude threshold such that any segment with at least one channel whose amplitude crosses the provided threshold will be marked for rejection. HAPPE+ER suggests an artifact threshold of -200 to 200 for infant data, and -150 to 150 for child, adolescent, and adult data. However, users are strongly encouraged to run the semi-automated HAPPE+ER setting on at least several files in their dataset to visually check that the selected amplitude results in appropriate segment rejection in their own datasets. Joint probability-based rejection catches other classes of artifacts, especially high-frequency artifacts like muscle artifact. Two joint probabilities are calculated with EEGLAB’s pop_jointprob function. The joint probability of an electrode’s activity in a segment given that same electrode’s activity in all other segments is calculated (single electrode probability), and the joint probability of an electrode’s activity in a segment given all other electrodes’ activities for that same segment is calculated (electrode group probability). These joint probabilities are evaluated such that any segment is marked for rejection when either (1) a channel’s single electrode probability or (2) its electrode group probability is outside of 3 standard deviations from the mean. All segments marked from the user-selected steps are then rejected simultaneously in a single step. Notably, this segment rejection step may be run on all user-specified channels, or on a subset of channels for a specific region of interest (ROI). The ROI-channel subset option allows users to tailor segment rejection for a specific ROI analysis and potentially retain more data per individual if that ROI is less artifact-contaminated relative to other ROIs in the channels selected for HAPPE+ER processing.

### Interpolation of Bad Channels

For all HAPPE+ER runs, regardless of segmentation options, any channels removed during the bad channel rejection processing step are now subject to spherical interpolation (with Legendre polynomials up to the 7th order) of their signal. Channel interpolation repopulates data for the complete channel set specified by the user and reduces bias in re-referencing if the average re-reference option is selected. The identity of all interpolated channels, if any, for a file are recorded in HAPPE’s processing report for users who wish to monitor the percentage or identity of interpolated channels in their datasets before further analysis.

### Re-Referencing (Optional)

HAPPE+ER offers users the choice to re-reference the EEG data. If re-referencing, the user may specify either re-referencing using an average across all channels (i.e., average re-reference) or using a channel subset of one or multiple channels. For both re-referencing options, only channels within the user-specified channel subset selected for HAPPE+ER processing can be used for re-referencing. Re-referencing also reduces artifact signals that exist consistently across electrodes, including residual line-noise. During re-referencing, if there is a prior reference channel (e.g., an online reference channel), that channel’s data is recovered and included in the re-referenced dataset. For example, EGI data is typically online-referenced to channel CZ. In this example, users could now recover data at channel CZ by re-referencing to any other channel or channels in this step.

### HAPPE+ER Outputs

HAPPE+ER generates several folders containing EEG files that are located within the user-specified folder of files for processing. EEG files are saved out after several intermediate processing steps so that users can explore in-depth and visualize how those steps affected the EEG signal in their own datasets. The intermediate files are separated into folders based on the level of processing performed on the data and include: 1) data after filtering to 100 Hz and line-noise reduction, 2) data post-bad channel rejection, 3) post-wavelet-thresholded data, and 4) data filtered at the user-specified ERP cutoffs. If segmenting is enabled, HAPPE+ER outputs one to three additional intermediate files: 5) post-segmented EEG data (always), 6) baseline-corrected data (if baseline correction is enabled), and 7) interpolated data (if bad data interpolation is enabled). If segment rejection is selected, HAPPE+ER saves the data post-segment rejection as well.

HAPPE+ER outputs fully processed files that are suitable inputs for further analyses in one of several formats, selected by the user at the start of the HAPPE+ER run, to increase compatibility with other software for data visualizations or statistical analyses. Options include .mat, .set, and .txt formats. We recommend using the .txt file format, which outputs three files in total: 1) A .txt file containing the average value across trials for each electrode at each sampling timepoint, 2) A .txt file containing the data for each electrode for each individual trial, and 3) An EEGLAB .set file of the fully processed EEG. Alongside the fully processed data, HAPPE+ER also outputs the HAPPE Data Quality Assessment Report and the HAPPE Pipeline Quality Assessment Report, each described in detail below, for the file batch. Finally, if the users ran HAPPE+ER in the semi-automated setting, the software generates an image for each file containing the fully processed data’s power spectrum.

### HAPPE Data Quality Assessment Report

HAPPE generates a report table of descriptive statistics and data metrics for each EEG file in the batch in a single spreadsheet to aid in quickly and effectively evaluating data quality across participants within or across studies. The report table with all these metrics is provided as a .csv file in the “quality_assessment_outputs” folder generated during HAPPE+ER. We describe each of these metrics below to facilitate their use to determine and report data quality.

#### File Length in Seconds

HAPPE+ER outputs the length, in seconds, of each file prior to processing.

#### Number of Segments Before Segment Rejection and Number of Segments Post Segment Rejection

HAPPE+ER reports the number of segments before segment rejection and post segment rejection. If segment rejection is not enabled, these numbers are identical. If the user enabled segment rejection in HAPPE+ER, they may evaluate the number of data segments remaining post-rejection for each file to identify any files that cannot contribute enough clean data to be included in further analyses (user discretion based on study design and ERP of interest). The user may also easily tabulate the descriptive statistics for remaining segments to report in their manuscript’s Methods section (e.g., the mean and standard deviation of the number of usable data segments per file in their study).

#### Percent Good Channels Selected and Interpolated Channel IDs

The percentage of channels contributing un-interpolated data (“good channels”) and the identity of interpolated channels are provided. Users wishing to limit the amount of interpolated data in further analyses can easily identify files for removal using these two metrics.

#### ICA-Related Metrics

As HAPPE+ER does not perform ICA on ERP data, the metrics measuring ICA performance in HAPPE 2.0 are assigned “NA.”

#### Channels Interpolated for Each Segment

If the user selected the Data Interpolation within Segments option of the additional segmenting options, HAPPE+ER will output a list of segments and the channels interpolated within each segment for each file. Otherwise, it will output “N/A.” Users wishing to limit the amount of interpolated data in further analyses can easily identify files for removal using this metric.

### HAPPE Pipeline Quality Assessment Report

For each run, HAPPE+ER additionally generates a report table of descriptive statistics and data metrics for each EEG file in the batch in a single spreadsheet to aid in quickly and effectively evaluating how well the pipeline performed across participants within or across studies. Note that these metrics may also be reported in manuscript methods sections as indicators of how data manipulations changed the signal during preprocessing. The report table with all these metrics is provided as a .csv file in the “quality_assessment_outputs” folder generated during HAPPE+ER processing.

#### r pre/post linenoise removal

HAPPE+ER automatically outputs cross-correlation values at and near the specified line noise frequency (correlation between data at each frequency before and after line noise removal). These cross-correlation values can be used to evaluate the performance of line noise removal, as the correlation pre- and post-line noise removal should be lower at the specified frequency, but not at the surrounding frequencies beyond 1 to 2 Hz. HAPPE+ER will automatically adjust which frequencies are reported depending on the user-identified line noise frequency. This metric can also be used to detect changes in how much line noise is present during the recordings (e.g. if generally cross-correlation values are high when study protocol is followed, indicating low line-noise removal from the data, but a staff member forgets to remove their cell phone from the recording booth for several testing sessions, the degree of line noise removal for those files summarized by this metric could be used as a flag to check in on site compliance with acquisition protocols).

#### r pre/post wav-threshold

HAPPE+ER automatically outputs the cross-correlation values before and after wavelet thresholding across all frequencies and specifically at 0.5 Hz, 1 Hz, 2 Hz, 5 Hz, 8 Hz, 12 Hz, 20 Hz, 30 Hz, 45 Hz, and 70 Hz. These specific frequencies were selected to cover all canonical frequency bands across the lifespan from delta through high-gamma as well as the low-frequencies retained in ERP analyses. These cross-correlation values can be used to evaluate the performance of waveleting on the data for each file. For example, if cross-correlation values are below 0.2 for all participants in the sample, the wavelet thresholding step has not progressed as intended (users are advised to first check their sampling rate in this case and visualize several raw data files). Note that this measure may also be used to exclude individual files from further analysis based on dramatic signal change during waveleting (indicating high degree of artifact), for example if the 0.5 Hz or all-data cross-correlations are below some threshold set by the user (e.g., 3 standard deviations from the median or mean, r values below 0.2)

Through these quality assessment reports, HAPPE+ER aims to provide a rich, quantifiable, yet easily accessible way to effectively evaluate data quality for even very large datasets in the context of automated processing. Visual examination of each file is not required, although it is available. Over and above the purposes of rejecting files that no longer meet quality standards for a study and evaluating HAPPE+ER performance on a given dataset, we also hope to encourage more rigorous reporting of data quality metrics in manuscripts by providing these outputs already tabulated and easily transformed into descriptive statistics for inclusion in reports. Users may also wish to include one or several of these metrics as continuous nuisance covariates in statistical analyses to better account for differences in data quality between files or verify whether there are statistically significant differences in data quality post-processing between study groups of interest.

Several metrics may also be useful in evaluating study progress remotely to efficiently track the integrity of the system and data collection protocols. For example, the r pre/post line noise removal metric may indicate environmental or protocol deviations that cause significant increases in line noise in the data, and the Percent Good Channels Selected and Interpolated Channel ID metrics can be used to track whether the net/cap is being applied and checked for signal quality prior to paradigms or whether a channel (or channels) is in need of repair. For example, if the T6 electrode starts consistently returning bad data for a specific net/cap, it may need to be examined for repair. For further guidance about using the processing report metrics to evaluate data, users may consult the User Guide distributed with HAPPE+ER software.

### Creating ERPs and Calculating ERP Values

HAPPE+ER comes with an optional post-processing script, called ‘generateERPs’, with the capability to generate ERP waveforms and perform a series of calculations on the resulting ERPs. This script is separate from the HAPPE+ER pipeline’s script to encourage users to check the quality of their data and HAPPE+ER’s performance prior to generating the ERP figures and measures. Any files that do not pass data quality thresholds should be removed from the outputs folder prior to running the generateERPs script, otherwise they will be included in the subsequent figures and metrics. Much like HAPPE+ER, generateERPs runs on input taken directly from the command line, continuing HAPPE+ER’s aim of making processing accessible to researchers of all levels of programming familiarity. The user must simply provide the full path of the processed folder created during the initial HAPPE+ER run and answer the prompts that follow.

To create ERPs, users can select their channels of interest in a nearly identical manner to channel selection in the HAPPE+ER pipeline (see above for more details). The only difference is that the 10-20 channels are not automatically included for the inclusion method of selecting channels. As part of channel selection, the user can additionally choose to include or exclude channels marked as “bad” during the HAPPE+ER processing run. If the user decides to exclude the bad channels detected during HAPPE+ER, they should first make a new .csv file that includes only the file names and bad channels columns from the Data Quality Assessment Report file generated by HAPPE+ER. This .csv file should be placed in the same folder as the processed data prior to running the generateERPs script. Note that if any files were removed post-processing due to insufficient data quality, they should be removed from the rows of this .csv as well.

The user is also asked whether they want to calculate a set of standard measures associated with the ERP for each file in the batch for subsequent statistical analysis. If so, the user must also specify: 1) latency windows of interest (e.g. 50-90 milliseconds post-stimulus), 2) whether they anticipate a maximum or a minimum to be present in that window (i.e. a positive or negative ERP component, respectively), and 3) whether to calculate area under the curve using temporal windows as bounds, using zero crossings present in the ERP data as bounds, or reporting measures with both methods (see Figure 5 and below).

**Figure 5.**
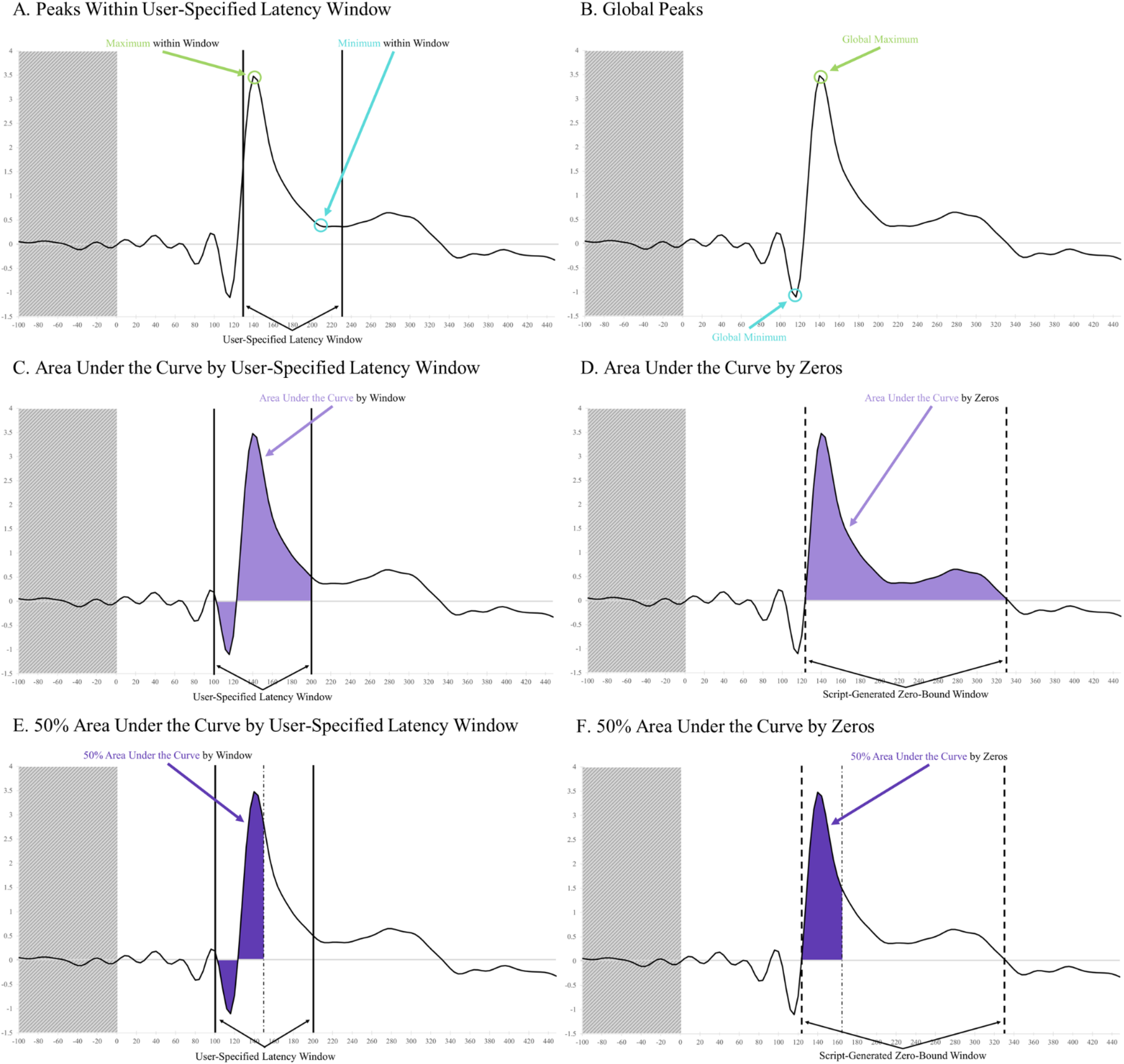
In all panels, the shaded area is not included in calculations. A) Circles the maximum (green) and minimum (blue) values within a specified latency window as indicated by vertical black lines. B) Circles the maximum (green) and minimum (blue) values across the entire ERP waveform. C) Area under the curve is represented in light purple, with user-specified boundaries indicated by solid black vertical lines. D) Area under the curve is represented in light purple, and the boundaries created by the script at zero crossings in the data as dashed lines. E) 50% area under the curve is represented in dark purple. User-specified boundaries are indicated by solid black vertical lines. The alternating dot-dash line represents the latency at which 50% area under the curve is reached within the window. F) 50% area under the curve is represented in dark purple. Script-generated boundaries at zero-crossings are indicated by dashed black vertical lines. The alternating dot-dash line represents the latency at which 50% area under the curve is reached within the window.

The generateERPs script will create an ERP timeseries for each subject as well as an average ERP timeseries across subjects, which are saved in a .csv output file. The name of this file is “allSubjects_generatedERPs” plus any suffix associated with the selected data. Three figures of these ERPs are also produced as generateERPs runs: 1) a figure containing the ERP of each subject, 2) the average ERP across subjects, and 3) a combination of the first two figures. A description of these figures is included in the MATLAB command line for the user’s convenience. If enabled, generateERPs calculates the following values for each file and the average ERP across files, outputting them in an additional .csv file.

#### Peak Amplitudes and Latencies

For each user-specified temporal window, generateERPs calculates the specified peak (either maximum or minimum depending on user input) and the latency at which it occurs (Figure 5). The user may specify the same temporal window twice to request both a maximum and minimum within that window. Additionally, the global maximum and minimum of the timeseries (Figure 5), a list of all maximums, a list of all minimums, and the latencies associated with each value are reported.

#### Area Under the Curve

If the user has selected area under the curve based on windows, generateERPs calculates the area under the curve using the user-specified latency windows’ start and end times as the upper and lower bounds (Figure 5). This method also reports the global area under the curve, calculated using absolute values, for the entire ERP waveform post-stimulus onset.

If the user has selected to calculate area under the curve based on zero crossings, generateERPs locates the zero crossings in the ERP and creates new latency windows using these crossing points, the starting latency, and the ending latency as bounds (Figure 5).

#### 50% Area Under the Curve and Latencies

If the user has selected 50% area under the curve based on windows, generateERPs calculates 50% of area under the curve for each user-specified latency window for bounds and the latency at which it is reached (Figure 5). This selection also reports the 50% area under the curve for the entire ERP waveform post-stimulus onset and its associated latency.

If the user has selected to calculate 50% area under the curve based on zeros, generateERPs creates latency windows using zero crossings in the ERP as bounds and reports the value and latency for this metric (Figure 5).

We provide this script with the hopes of facilitating ERP visualization and analysis.

Additionally, the speed, automation, and inherent compatibility with HAPPE+ER aligns directly with HAPPE+ER’s goals of providing an accessible and standardized method of examining EEG/ERP data.

### Implementing HAPPE+ER

HAPPE+ER runs entirely through the MATLAB command line, collecting processing parameters without the user needing to navigate or alter the pipeline’s code. This reduces the chance of accidentally breaking the code, entering incorrect parameters for the desired analysis, and the need to have prior knowledge of coding or MATLAB, enabling users of wide range of backgrounds and levels of familiarity to easily run the pipeline. To run HAPPE+ER, simply open MATLAB, navigate to the HAPPE 2.0 folder, and open the HAPPE 2.0 script. In the “Editor” tab at the top of the screen, click “Run” and follow the prompts as they appear in the MATLAB command line. After entering all relevant inputs to the command line, HAPPE+ER will run automatically through completion.

HAPPE+ER code and user guide are freely available at: https://github.com/PINE-Lab/HAPPE.

## Appendix

Key for reading the repeated-measures ANOVA results:

| Sample | Column 1 | Column 2 | Column 3 | Column 4 |
| --- | --- | --- | --- | --- |
| Row 1 | < | > |  |  |
|  | Row 1 < Column 1 * | Row 1 > Column 2 * | Row 1 = Column 3 | Not Examined |

\* $p < 0.05$

Complete pattern results for the ANOVAs described in the above paper.

### Wavelet Level Optimization

#### Adults

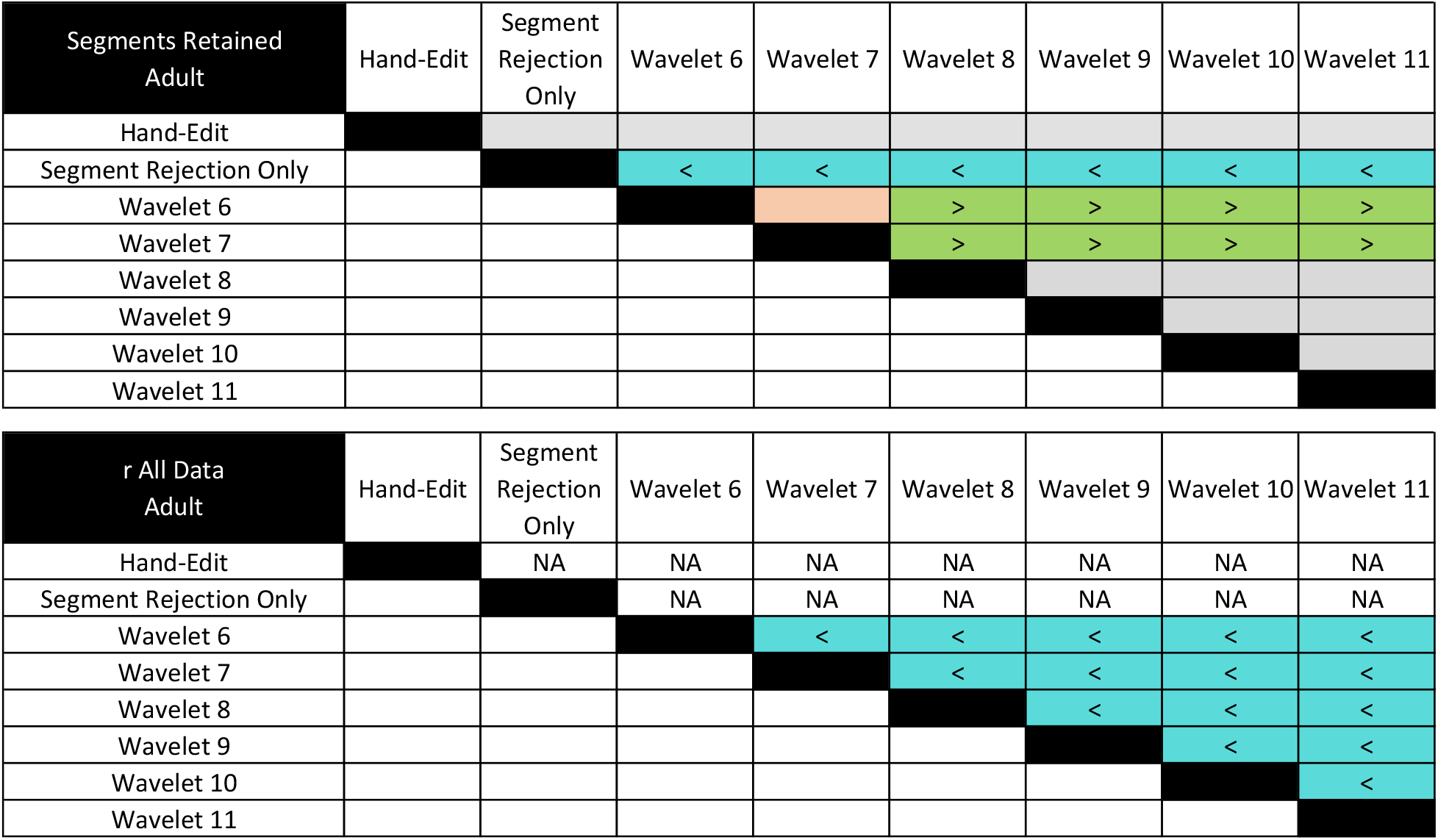

#### 4-Month

| Segments Retained 4-Month | Hand-Edit | Segment Rejection Only | Wavelet 8 | Wavelet 9 | Wavelet 10 | Wavelet 11 | Wavelet 12 | Wavelet 13 |
| --- | --- | --- | --- | --- | --- | --- | --- | --- |
| Hand-Edit |  | < | < | < | < | < | < | < |
| Segment Rejection Only |  |  | < | < | < | < | < | < |
| Wavelet 8 |  |  |  |  |  |  | > | > |
| Wavelet 9 |  |  |  |  |  |  | > | > |
| Wavelet 10 |  |  |  |  |  |  |  |  |
| Wavelet 11 |  |  |  |  |  |  |  |  |
| Wavelet 12 |  |  |  |  |  |  |  |  |
| Wavelet 13 |  |  |  |  |  |  |  |  |

| r All Data<br>4-Month | Hand-Edit | Segment<br>Rejection<br>Only | Wavelet 8 | Wavelet 9 | Wavelet 10 | Wavelet 11 | Wavelet 12 | Wavelet 13 |
| --- | --- | --- | --- | --- | --- | --- | --- | --- |
| Hand-Edit |  | NA | NA | NA | NA | NA | NA | NA |
| Segment Rejection Only |  |  | NA | NA | NA | NA | NA | NA |
| Wavelet 8 |  |  |  | < | < | < | < | < |
| Wavelet 9 |  |  |  |  | < | < | < | < |
| Wavelet 10 |  |  |  |  |  | < | < | < |
| Wavelet 11 |  |  |  |  |  |  | < | < |
| Wavelet 12 |  |  |  |  |  |  |  | < |
| Wavelet 13 |  |  |  |  |  |  |  |  |

#### 10-Month

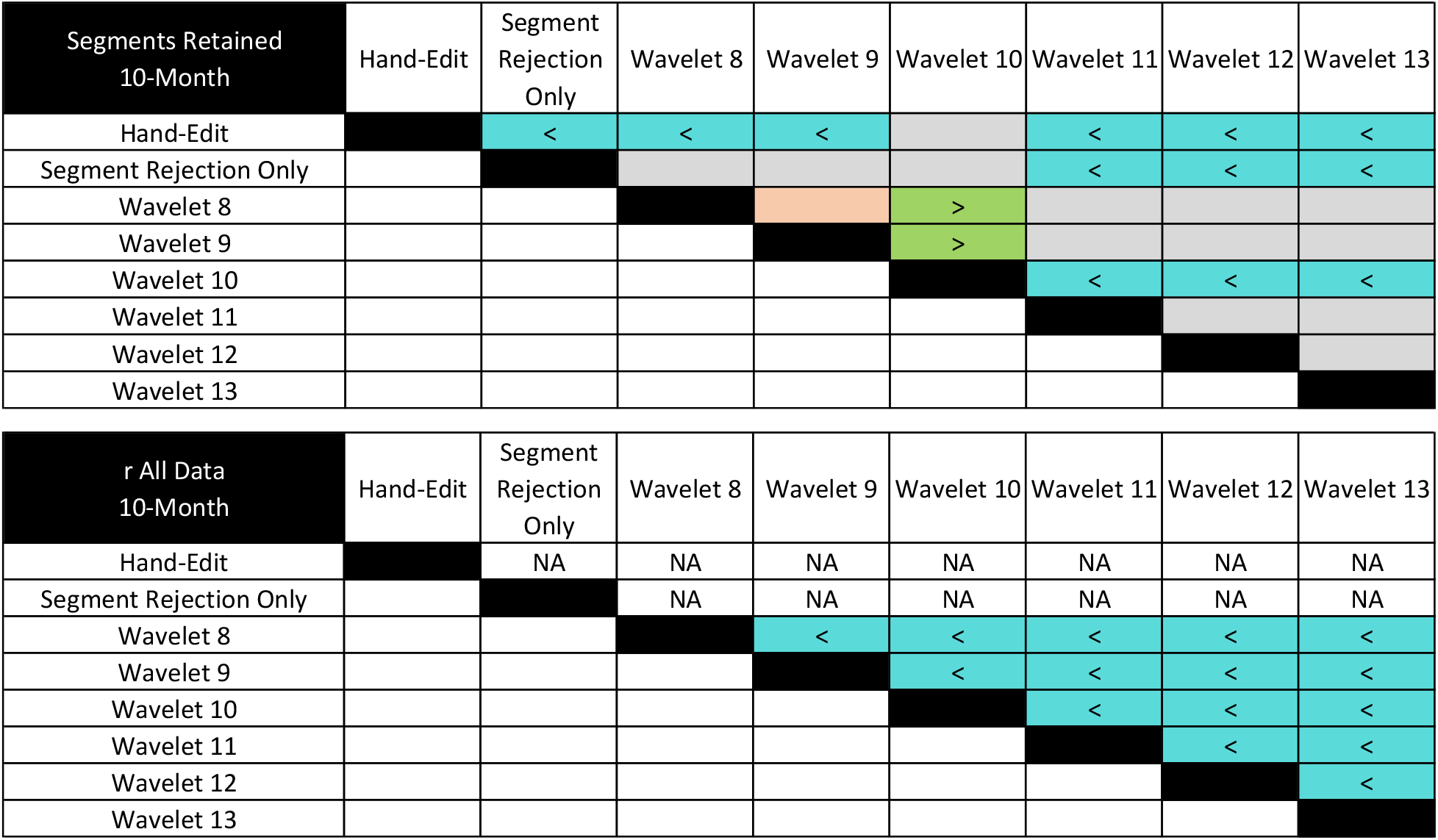

### Artifact Rejection Methods

#### Adults

Artifact Rejection Methods Adults
| Segments Retained<br>Adult | Hand-Edit | Segment<br>Rejection<br>Only | ICA Only | Waveleting<br>Only | Waveleting<br>+ ICA | wICA Only | wICA + ICA |
| --- | --- | --- | --- | --- | --- | --- | --- |
| Hand-Edit |  |  |  |  |  |  |  |
| Segment Rejection Only |  |  | < |  | < |  |  |
| ICA Only |  |  |  |  |  | > | > |
| Waveleting Only |  |  |  |  |  | > |  |
| Waveleting + ICA |  |  |  |  |  | > |  |
| wICA Only |  |  |  |  |  |  | < |
| wICA + ICA |  |  |  |  |  |  |  |

| r All Data Adult | Hand-Edit | Segment Rejection Only | ICA Only | Waveleting Only | Waveleting + ICA | wICA Only | wICA + ICA |
| --- | --- | --- | --- | --- | --- | --- | --- |
| Hand-Edit |  | NA | NA | NA | NA | NA | NA |
| Segment Rejection Only |  |  | NA | NA | NA | NA | NA |
| ICA Only |  |  |  | < |  | < |  |
| Waveleting Only |  |  |  |  | > |  | > |
| Waveleting + ICA |  |  |  |  |  | < |  |
| wICA Only |  |  |  |  |  |  | > |
| wICA + ICA |  |  |  |  |  |  |  |

| N1 Amplitude Adult | Hand-Edit | Segment Rejection Only | ICA Only | Waveleting Only | Waveleting + ICA | wICA Only | wICA + ICA |
| --- | --- | --- | --- | --- | --- | --- | --- |
| Hand-Edit |  |  |  | > |  |  |  |
| Segment Rejection Only |  |  |  |  |  |  |  |
| ICA Only |  |  |  | > |  |  |  |
| Waveleting Only |  |  |  |  | < |  | < |
| Waveleting + ICA |  |  |  |  |  |  |  |
| wICA Only |  |  |  |  |  |  |  |
| wICA + ICA |  |  |  |  |  |  |  |

| N1 to P1 Peak-to-Peak Adult | Hand-Edit | Segment Rejection Only | ICA Only | Waveleting Only | Waveleting + ICA | wICA Only | wICA + ICA |
| --- | --- | --- | --- | --- | --- | --- | --- |
| Hand-Edit |  |  | > |  | > |  | > |
| Segment Rejection Only |  |  | > |  | > |  | > |
| ICA Only |  |  |  | < |  | < |  |
| Waveleting Only |  |  |  |  | > |  | > |
| Waveleting + ICA |  |  |  |  |  | < | < |
| wICA Only |  |  |  |  |  |  | > |
| wICA + ICA |  |  |  |  |  |  |  |

| P1 to N2 Peak-to-Peak Adult | Hand-Edit | Segment Rejection Only | ICA Only | Waveleting Only | Waveleting + ICA | wICA Only | wICA + ICA |
| --- | --- | --- | --- | --- | --- | --- | --- |
| Hand-Edit |  |  | > |  | > |  | > |
| Segment Rejection Only |  |  | > |  | > |  | > |
| ICA Only |  |  |  |  | > | < |  |
| Waveleting Only |  |  |  |  | > | < | > |
| Waveleting + ICA |  |  |  |  |  | < | < |
| wICA Only |  |  |  |  |  |  | > |
| wICA + ICA |  |  |  |  |  |  |  |

#### 4-Month

##### MARA 0.5

4-Month MARA 0.5
| Segments Retained 4-Month | Hand-Edit | Segment Rejection Only | ICA Only | Waveleting Only | Waveleting + ICA | wICA Only | wICA + ICA |
| --- | --- | --- | --- | --- | --- | --- | --- |
| Hand-Edit |  | < | > | < | > | > |  |
| Segment Rejection Only |  |  | > | < | > | > | > |
| ICA Only |  |  |  | < |  | < | < |
| Waveleting Only |  |  |  |  | > | > | > |
| Waveleting + ICA |  |  |  |  |  | < | < |
| wICA Only |  |  |  |  |  |  |  |
| wICA + ICA |  |  |  |  |  |  |  |

| r All Data by 4-Month | Hand-Edit | Segment Rejection Only | ICA Only | Waveleting Only | Waveleting + ICA | wICA Only | wICA + ICA |
| --- | --- | --- | --- | --- | --- | --- | --- |
| Hand-Edit |  | NA | NA | NA | NA | NA | NA |
| Segment Rejection Only |  |  | NA | NA | NA | NA | NA |
| ICA Only |  |  |  | < |  | < | < |
| Waveleting Only |  |  |  |  | > |  | > |
| Waveleting + ICA |  |  |  |  |  | < |  |
| wICA Only |  |  |  |  |  |  | > |
| wICA + ICA |  |  |  |  |  |  |  |

##### iMARA 0.2

| Segments Retained 4-Month | Hand-Edit | Segment Rejection Only | ICA Only | Waveleting Only | Waveleting + ICA | wICA Only | wICA + ICA |
| --- | --- | --- | --- | --- | --- | --- | --- |
| Hand-Edit |  | < | < | < | < | > | < |
| Segment Rejection Only |  |  | < | < | < | > |  |
| ICA Only |  |  |  | < |  | > | > |
| Waveleting Only |  |  |  |  |  | > | > |
| Waveleting + ICA |  |  |  |  |  | > | > |
| wICA Only |  |  |  |  |  |  | < |
| wICA + ICA |  |  |  |  |  |  |  |

| r All Data 4-Month | Hand-Edit | Segment Rejection Only | ICA Only | Waveleting Only | Waveleting + ICA | wICA Only | wICA + ICA |
| --- | --- | --- | --- | --- | --- | --- | --- |
| Hand-Edit |  | NA | NA | NA | NA | NA | NA |
| Segment Rejection Only |  |  | NA | NA | NA | NA | NA |
| ICA Only |  |  |  | < |  | < |  |
| Waveleting Only |  |  |  |  | > |  | > |
| Waveleting + ICA |  |  |  |  |  | < | < |
| wICA Only |  |  |  |  |  |  | > |
| wICA + ICA |  |  |  |  |  |  |  |

| N1 Amplitude<br>4-Month | Hand-Edit | Segment<br>Rejection<br>Only | ICA Only | Waveleting<br>Only | Waveleting<br>+ ICA | wICA Only | wICA + ICA |
| --- | --- | --- | --- | --- | --- | --- | --- |
| Hand-Edit |  |  | < |  | < |  | < |
| Segment Rejection Only |  |  | < |  | < |  |  |
| ICA Only |  |  |  |  |  | > |  |
| Waveleting Only |  |  |  |  | < |  |  |
| Waveleting + ICA |  |  |  |  |  | > |  |
| wICA Only |  |  |  |  |  |  |  |
| wICA + ICA |  |  |  |  |  |  |  |

| N1 to P1 Peak-to-Peak<br>4-Month | Hand-Edit | Segment<br>Rejection<br>Only | ICA Only | Waveleting<br>Only | Waveleting<br>+ ICA | wICA Only | wICA + ICA |
| --- | --- | --- | --- | --- | --- | --- | --- |
| Hand-Edit |  |  | > | > | > |  | > |
| Segment Rejection Only |  |  | > |  | > |  |  |
| ICA Only |  |  |  | < |  | < | < |
| Waveleting Only |  |  |  |  | > | < |  |
| Waveleting + ICA |  |  |  |  |  | < | < |
| wICA Only |  |  |  |  |  |  | > |
| wICA + ICA |  |  |  |  |  |  |  |

| P1 to N2 Peak-to-Peak<br>4-Month | Hand-Edit | Segment<br>Rejection<br>Only | ICA Only | Waveleting<br>Only | Waveleting<br>+ ICA | wICA Only | wICA + ICA |
| --- | --- | --- | --- | --- | --- | --- | --- |
| Hand-Edit |  |  | > |  | > | < | > |
| Segment Rejection Only |  |  | > |  | > | < |  |
| ICA Only |  |  |  | < |  | < | < |
| Waveleting Only |  |  |  |  |  | < |  |
| Waveleting + ICA |  |  |  |  |  | < |  |
| wICA Only |  |  |  |  |  |  | > |
| wICA + ICA |  |  |  |  |  |  |  |

#### 10-Month

##### MARA0.5

10-Month MARA 0.5
| Segments Retained<br>10-Month | Hand-Edit | Segment<br>Rejection<br>Only | ICA Only | Waveleting<br>Only | Waveleting<br>+ ICA | wICA Only | wICA + ICA |
| --- | --- | --- | --- | --- | --- | --- | --- |
| Hand-Edit |  | < |  | < |  |  | < |
| Segment Rejection Only |  |  |  |  |  | > | < |
| ICA Only |  |  |  |  |  |  |  |
| Waveleting Only |  |  |  |  |  | > | < |
| Waveleting + ICA |  |  |  |  |  |  |  |
| wICA Only |  |  |  |  |  |  | < |
| wICA + ICA |  |  |  |  |  |  |  |

| r All Data<br>10-Month | Hand-Edit | Segment<br>Rejection<br>Only | ICA Only | Waveleting<br>Only | Waveleting<br>+ ICA | wICA Only | wICA + ICA |
| --- | --- | --- | --- | --- | --- | --- | --- |
| Hand-Edit |  | NA | NA | NA | NA | NA | NA |
| Segment Rejection Only |  |  | NA | NA | NA | NA | NA |
| ICA Only |  |  |  | < | < | < | < |
| Waveleting Only |  |  |  |  | > | < | > |
| Waveleting + ICA |  |  |  |  |  | < | < |
| wICA Only |  |  |  |  |  |  | > |
| wICA + ICA |  |  |  |  |  |  |  |

##### iMARA 0.2

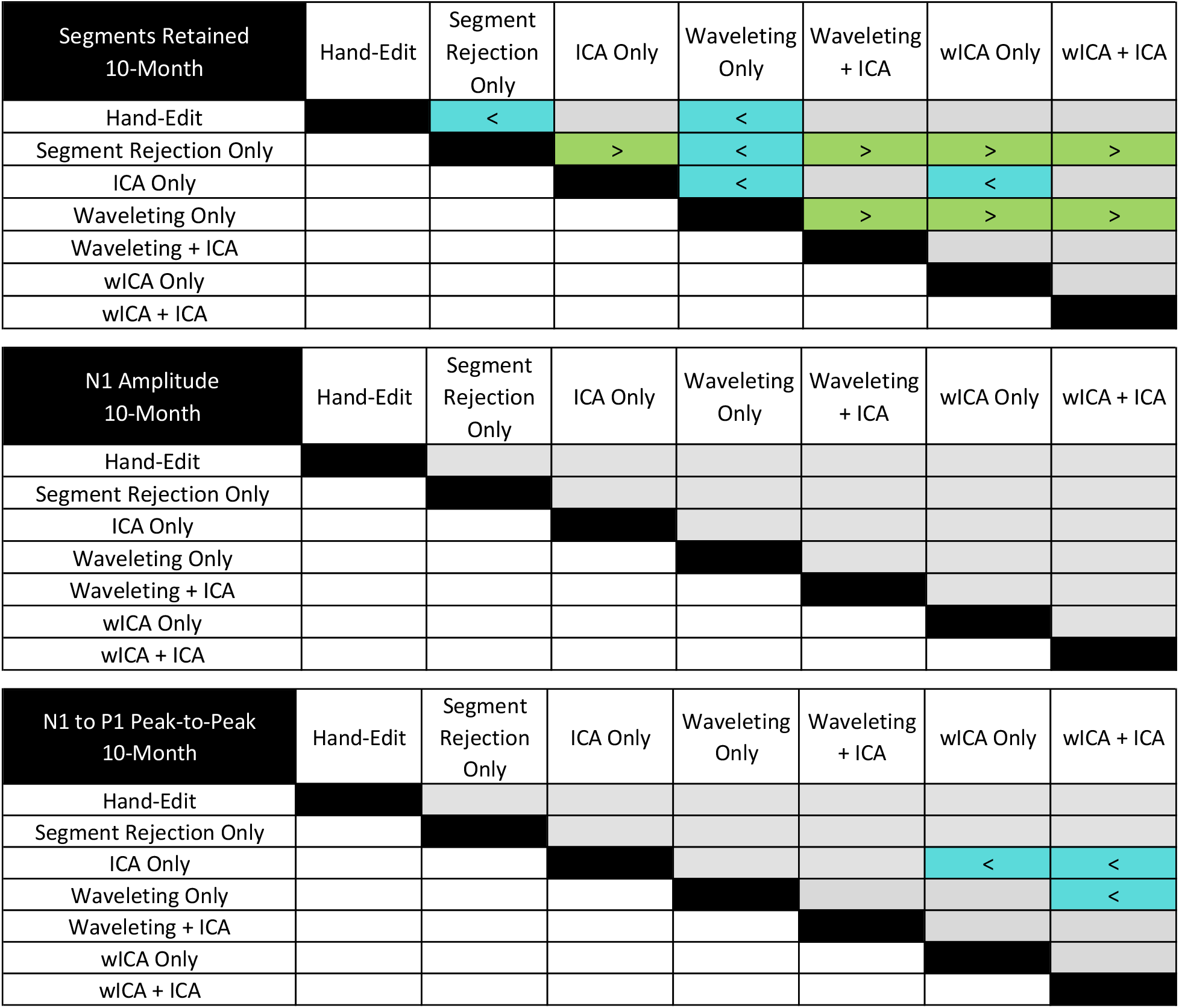
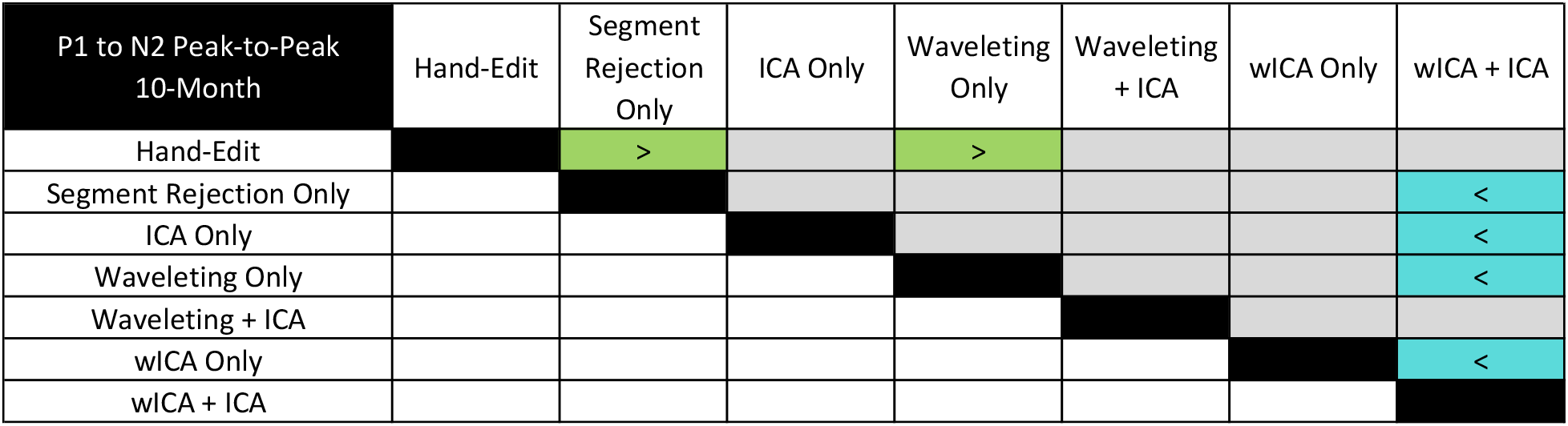

#### ICA Optimization

| Segments Retained<br>ICA Optimization (4-Month) | Hand-Edit | Nothing | Waveleting + ICA<br>using iMARA (0.5) | Waveleting + ICA<br>using iMARA (0.2) | Waveleting + ICA<br>using ICLabel (0.5) | Waveleting + ICA<br>using ICLabel (0.8) | Waveleting + ICA<br>using adjusted-ADJUST |
| --- | --- | --- | --- | --- | --- | --- | --- |
| Hand-Edit |  | < | < | < | < |  |  |
| Nothing |  |  | < | < | < |  | > |
| Waveleting + ICA using iMARA (0.5) |  |  |  | > |  | > | > |
| Waveleting + ICA using iMARA (0.2) |  |  |  |  | < | > | > |
| Waveleting + ICA using ICLabel (0.5) |  |  |  |  |  | > | > |
| Waveleting + ICA using ICLabel (0.8) |  |  |  |  |  |  |  |
| Waveleting + ICA using adjusted-ADJUST |  |  |  |  |  |  |  |

| N1 Amplitude<br>ICA Optimization (4-Month) | Hand-Edit | Nothing | Waveleting + ICA<br>using iMARA (0.5) | Waveleting + ICA<br>using iMARA (0.2) | Waveleting + ICA<br>using ICLabel (0.5) | Waveleting + ICA<br>using ICLabel (0.8) |
| --- | --- | --- | --- | --- | --- | --- |
| Hand-Edit |  |  | < | < | < | < |
| Nothing |  |  | < | < | < | < |
| Waveleting + ICA using iMARA (0.5) |  |  |  | > |  | > |
| Waveleting + ICA using iMARA (0.2) |  |  |  |  | < |  |
| Waveleting + ICA using ICLabel (0.5) |  |  |  |  |  | > |
| Waveleting + ICA using ICLabel (0.8) |  |  |  |  |  |  |

| N1 to P1 Peak-to-Peak<br>ICA Optimization (4-Month) | Hand-Edit | Nothing | Waveleting + ICA<br>using iMARA (0.5) | Waveleting + ICA<br>using iMARA (0.2) | Waveleting + ICA<br>using ICLabel (0.5) | Waveleting + ICA<br>using ICLabel (0.8) |
| --- | --- | --- | --- | --- | --- | --- |
| Hand-Edit |  |  | > | > | > | > |
| Nothing |  |  | > | > | > | > |
| Waveleting + ICA using iMARA (0.5) |  |  |  | < |  | < |
| Waveleting + ICA using iMARA (0.2) |  |  |  |  | > |  |
| Waveleting + ICA using ICLabel (0.5) |  |  |  |  |  | < |
| Waveleting + ICA using ICLabel (0.8) |  |  |  |  |  |  |

| P1 to N2 Peak-to-Peak<br>ICA Optimization (4-Month) | Hand-Edit | Nothing | Waveleting + ICA<br>using iMARA (0.5) | Waveleting + ICA<br>using iMARA (0.2) | Waveleting + ICA<br>using ICLabel (0.5) | Waveleting + ICA<br>using ICLabel (0.8) |
| --- | --- | --- | --- | --- | --- | --- |
| Hand-Edit |  |  | > | > | > | > |
| Nothing |  |  | > | > | > | > |
| Waveleting + ICA using iMARA (0.5) |  |  |  | < |  | < |
| Waveleting + ICA using iMARA (0.2) |  |  |  |  | > |  |
| Waveleting + ICA using ICLabel (0.5) |  |  |  |  |  | < |
| Waveleting + ICA using ICLabel (0.8) |  |  |  |  |  |  |

## References

Bigdely-Shamlo, N., Mullen, T., Kothe, C., Su, K.M., Robbins, K.A., 2015. The PREP pipeline: Standardized preprocessing for large-scale EEG analysis. Front. Neuroinform. 9, 1–19. https://doi.org/10.3389/fninf.2015.00016

Cassani, R., Falk, T.H., Fraga, F.J., Cecchi, M., Moore, D.K., Anghinah, R., 2017. Towards automated electroencephalography-based Alzheimer’s disease diagnosis using portable low-density devices. Biomed. Signal Process. Control 33, 261–271. https://doi.org/10.1016/j.bspc.2016.12.009

Castellanos, N.P., Makarov, V.A., 2006. Recovering EEG brain signals: Artifact suppression with wavelet enhanced independent component analysis. J. Neurosci. Methods 158, 300–312. https://doi.org/10.1016/j.jneumeth.2006.05.033

da Cruz, J.R., Chicherov, V., Herzog, M.H., Figueiredo, P., 2018. An automatic pre-processing pipeline for EEG analysis (APP) based on robust statistics. Clin. Neurophysiol. 129, 1427– 1437. https://doi.org/10.1016/j.clinph.2018.04.600

Debnath, R., Buzzell, G.A., Morales, S., Bowers, M.E., Leach, S.C., Fox, N.A., 2020. The Maryland analysis of developmental EEG (MADE) pipeline. Psychophysiology 57, e13580. https://doi.org/10.1111/psyp.13580

Delorme, A., Makeig, S., 2004. EEGLAB: An open source toolbox for analysis of single-trial EEG dynamics including independent component analysis. J. Neurosci. Methods 134, 9–21. https://doi.org/10.1016/j.jneumeth.2003.10.009

Desjardins, J.A., van Noordt, S., Huberty, S., Segalowitz, S.J., Elsabbagh, M., 2021. EEG Integrated Platform Lossless (EEG-IP-L) pre-processing pipeline for objective signal quality assessment incorporating data annotation and blind source separation. J. Neurosci. Methods 347, 108961. https://doi.org/10.1016/j.jneumeth.2020.108961

Gabard-Durnam, L.J., Leal, A.S.M., Wilkinson, C.L., Levin, A.R., 2018. The harvard automated processing pipeline for electroencephalography (HAPPE): Standardized processing software for developmental and high-artifact data. Front. Neurosci. 12, 97. https://doi.org/10.3389/fnins.2018.00097

Gramfort, A., Luessi, M., Larson, E., Engemann, D.A., Strohmeier, D., Brodbeck, C., Parkkonen, L., Hämäläinen, M.S., 2014. MNE software for processing MEG and EEG data. Neuroimage 86, 446–460. https://doi.org/10.1016/j.neuroimage.2013.10.027

Haresign, M., Phillips, E., Whitehorn, M., Noreika, V., Jones, E.J.H., Leong, V., Wass, S. V, 2021. Automatic classification of ICA components from infant EEG using MARA. Biorxiv 1–43.

Hatz, F., Hardmeier, M., Bousleiman, H., Rüegg, S., Schindler, C., Fuhr, P., 2015. Reliability of fully automated versus visually controlled pre- and post-processing of resting-state EEG. Clin. Neurophysiol. 126, 268–274. https://doi.org/10.1016/j.clinph.2014.05.014

Lawhern, V., Hairston, W.D., Robbins, K., 2013. DETECT: A MATLAB Toolbox for Event Detection and Identification in Time Series, with Applications to Artifact Detection in EEG Signals. PLoS One 8, 62944. https://doi.org/10.1371/journal.pone.0062944

Leach, S.C., Morales, S., Bowers, M.E., Buzzell, G.A., Debnath, R., Beall, D., Fox, N.A., 2020. Adjusting ADJUST: Optimizing the ADJUST algorithm for pediatric data using geodesic nets. Psychophysiology 57, e13566. https://doi.org/10.1111/psyp.13566

Luck, S.J., 2014. An Introduction to the Event-Related Potential Technique, 2nd Revised., 2nd ed, The MIT press. MIT Press.

Mitra, P.P., Pesaran, B., 1999. Analysis of dynamic brain imaging data. Biophys. J. 76, 691–708. https://doi.org/10.1016/S0006-3495(99)77236-X

Mognon, A., Jovicich, J., Bruzzone, L., Buiatti, M., 2011. ADJUST: An automatic EEG artifact detector based on the joint use of spatial and temporal features. Psychophysiology 48, 229–240. https://doi.org/10.1111/j.1469-8986.2010.01061.x

Mullen, T., 2012. CleanLine EEGLAB Plugin.

Nolan, H., Whelan, R., Reilly, R.B., 2010. FASTER: Fully Automated Statistical Thresholding for EEG artifact Rejection. J. Neurosci. Methods 192, 152–162. https://doi.org/10.1016/j.jneumeth.2010.07.015

Oostenveld, R., Fries, P., Maris, E., Schoffelen, J.M., 2011. FieldTrip: Open source software for advanced analysis of MEG, EEG, and invasive electrophysiological data. Comput. Intell. Neurosci. 2011. https://doi.org/10.1155/2011/156869

Pedroni, A., Bahreini, A., Langer, N., 2019. Automagic: Standardized preprocessing of big EEG data. Neuroimage 200, 460–473. https://doi.org/10.1016/j.neuroimage.2019.06.046

Pion-Tonachini, L., Makeig, S., Kreutz-Delgado, K., 2017. Crowd labeling latent Dirichlet allocation. Knowl. Inf. Syst. 53, 749–765. https://doi.org/10.1007/s10115-017-1053-1

Sokol, S., 1976. Visually evoked potentials: Theory, techniques and clinical applications. Surv. Ophthalmol. https://doi.org/10.1016/0039-6257(76)90046-1

Tadel, F., Baillet, S., Mosher, J.C., Pantazis, D., Leahy, R.M., 2011. Brainstorm: A user-friendly application for MEG/EEG analysis. Comput. Intell. Neurosci. 2011. https://doi.org/10.1155/2011/879716

Varcin, K.J., Nelson, C.A., 2016. A developmental neuroscience approach to the search for biomarkers in autism spectrum disorder. Curr. Opin. Neurol. https://doi.org/10.1097/WCO.0000000000000298

Winkler, I., Debener, S., Muller, K.R., Tangermann, M., 2015. On the influence of high-pass filtering on ICA-based artifact reduction in EEG-ERP, in: Proceedings of the Annual International Conference of the IEEE Engineering in Medicine and Biology Society, EMBS. Institute of Electrical and Electronics Engineers Inc., pp. 4101–4105. https://doi.org/10.1109/EMBC.2015.7319296

Winkler, I., Haufe, S., Tangermann, M., 2011. Automatic Classification of Artifactual ICA-Components for Artifact Removal in EEG Signals. Behav. Brain Funct. 7, 1–15. https://doi.org/10.1186/1744-9081-7-30

